# Human reference gut microbiome comprising 5,414 prokaryotic species, including newly assembled genomes from under-represented Asian metagenomes

**DOI:** 10.1101/2020.11.09.375873

**Authors:** Chan Yeong Kim, Muyoung Lee, Sunmo Yang, Kyungnam Kim, Dongeun Yong, Hye Ryun Kim, Insuk Lee

## Abstract

Metagenome sampling bias for geographical location and lifestyle is partially responsible for the incomplete catalog of reference genomes of gut microbial species. Here, we present a substantially expanded microbiome catalog, the Human Reference Gut Microbiome (HRGM). Incorporating newly assembled 29,082 genomes from 845 fecal samples collected from three under-represented Asian countries—Korea, India, and Japan—the HRGM contains 232,098 non-redundant genomes of 5,414 representative prokaryotic species, >103 million unique proteins, and >274 million single-nucleotide variants. This is an over 10% increase from the largest reference database. The newly assembled genomes were enriched for members of the *Bacteroidaceae* family, including species associated with high-fiber and seaweed-rich diet. Single-nucleotide variant density was positively associated with the speciation rate of gut commensals. Ultra-deep sequencing facilitated the assembly of genomes of low-abundance taxa, and deep sequencing (>20 million read pairs) was needed for the profiling of low-abundance taxa. Importantly, the HRGM greatly improved the taxonomic and functional classification of sequencing reads from fecal samples. Finally, mapping homologous sequences for human auto-antigens onto the HRGM genomes revealed the association of commensal bacteria with high cross-reactivity potential with autoimmunity. The HRGM (www.mbiomenet.org/HRGM/) will facilitate the identification and functional analysis of disease-associated gut microbiota.

## Introduction

Human gut microbiome is considered the “second human genome” and plays a crucial role in various diseases^1,2^. Therefore, targeting gut microbes and their functional elements may provide novel therapeutic opportunities. The assembly of human reference genome, together with a catalog of protein-coding genes and genomic variants, led us to the era of genomic medicine. Likewise, transformation of human medicine by harnessing the gut microbes requires the cataloging of reference microbial genomes and their encoded functional elements. Conventional approaches for microbial genome assembly require microbial isolation and culture. Indeed, with the development of culturomics technology, the number of culturable gut microbes has increased greatly^3–6^. However, the culturable taxa are biased toward specific clades, and a large portion of the human gut microbiome remains unculturable^7–9^. To address this, culture-independent methods of metagenome assembly from whole-metagenomic shotgun sequencing (WMS) data have been developed.

Recently, three independent studies have consecutively released large collections of prokaryotic genomes, including many based on metagenome assembly^8–10^. The metagenome-assembled genomes (MAGs) from these studies were then combined with the genomic information deposited in public databases to generate integrated catalogs of prokaryotic genomes and proteins in the human gut^11^, the Unified Human Gastrointestinal Genome (UHGG) and Unified Human Gastrointestinal Protein (UHGP) catalogs, respectively. The UHGG contains 204,938 non-redundant genomes that represent 4,644 prokaryotic species and the UHGP catalogs approximately 95 million unique proteins.

Despite the latest advances, the current human gut microbiome catalog is incomplete, partially because the metagenome sampling is biased for geographical location and lifestyle. Specifically, the UHGG is strongly biased towards fecal samples collected in China, Denmark, Spain, and the US. In the present study, to account for the under-sampling of certain metagenomes, we assembled genomes from fecal samples collected from Korea, India, and Japan. Since the genome assembly of low-abundance species in most human fecal samples may require a much deeper sequencing than usually employed, we performed ultra-deep WMS (>30 Gbp or >100 million read pairs) of 90 fecal samples collected from Korea. We then collected public WMS data for 110 fecal samples from India and 805 fecal samples from Japan. We consequently assembled 29,082 prokaryotic genomes, and combined them with the UHGG genomes to generate the Human Reference Gut Microbiome (HRGM), which substantially expands the list of representative species, genomes, proteins, and single-nucleotide variants (SNVs) in the human gut microbiome. The HRGM is a freely available resource and will be invaluable to therapeutic targeting of the gut microbiota.

## Results

### Assembly of gut microbial genomes from Korea, India, and Japan

We assembled prokaryotic genomes using an in-house bioinformatics pipeline (**Supplementary Fig. 1a, Methods**), which is more exhaustive than similar approaches^8–11^ (**Supplementary Table 1**). For instance, we adopted an ensemble method for binning assembled contigs, as it showed better performance than individual binning tools^12,13^. We hypothesized that metagenomes harbored by individuals from under-represented geographical locations and lifestyles would expand the current catalog of human gut microbiome. Therefore, we performed *de novo* genome assembly of fecal samples from three Asian countries: Korea, India, and Japan (referred to here as KIJ samples, **Supplementary Table 2**). At the start of the current study, WMS data for 805 and 110 fecal samples from Japan and India, respectively, were publicly available but not included in the UHGG^14,15^. To complement these data, we generated WMS data for fecal samples collected from 90 donors recruited in Korea. We set the minimum completeness at 50% and the maximum contamination at 5% for genomes of minimum quality. We divided the genome bins into two groups: high quality (HQ) genomes with ≥90% completeness and ≤5% contamination, and medium quality (MQ) genomes (the remaining genomes). This yielded 29,082 KIJ sample MAGs: 7,767 from Korea, 563 from India, and 20,752 from Japan.

### Ultra-deep sequencing facilitates the genomic assembly of low-abundance taxa

To investigate the impact of metagenome sequencing depth on *de novo* genome assembly, we performed ultra-deep sequencing of the 90 Korean fecal samples (>30 Gbp or >100 million read pairs); the depth was approximately 5-fold deeper than the normal sequencing depth (**Fig. 1a**). Despite sequencing at the normal depth, fecal samples from Japan had a larger total read length than Korean samples because of a much larger sample size (**Fig. 1b**). For nine of the 90 Korean samples, approximately 60 Gbp was sequenced for the study of sequencing depth effect on genome assembly. We then generated 81 simulated WMS datasets (9 different depths for each of the 9 original samples with ~60 Gbp depth) and used the same pipeline of *de novo* genome assembly for all samples. As expected, the number of HQ and MQ genomes increased with the increasing sequencing depth. However, the growth rate simultaneously decreased and the proportion of HQ genomes became stable after the initial phase of rapid growth (**Fig. 1c**). Next, we investigated whether the increased sequencing depth improved the quality of assembled genomes. We compared the assembly quality of MAGs for the same species in two different simulated samples at adjacent sequencing depths (**Supplementary Fig. 2**; **Methods**). The quality of MAGs from the greater sequencing depth was significantly higher than that of genomes from the lower sequencing depth in terms of completeness, contamination, N50, and genome size (**Fig. 1d,e; Supplementary Fig. 3a,b**). However, the degree of improvement of the assembly quality diminished as the sequencing depth increased.

**Fig. 1.**
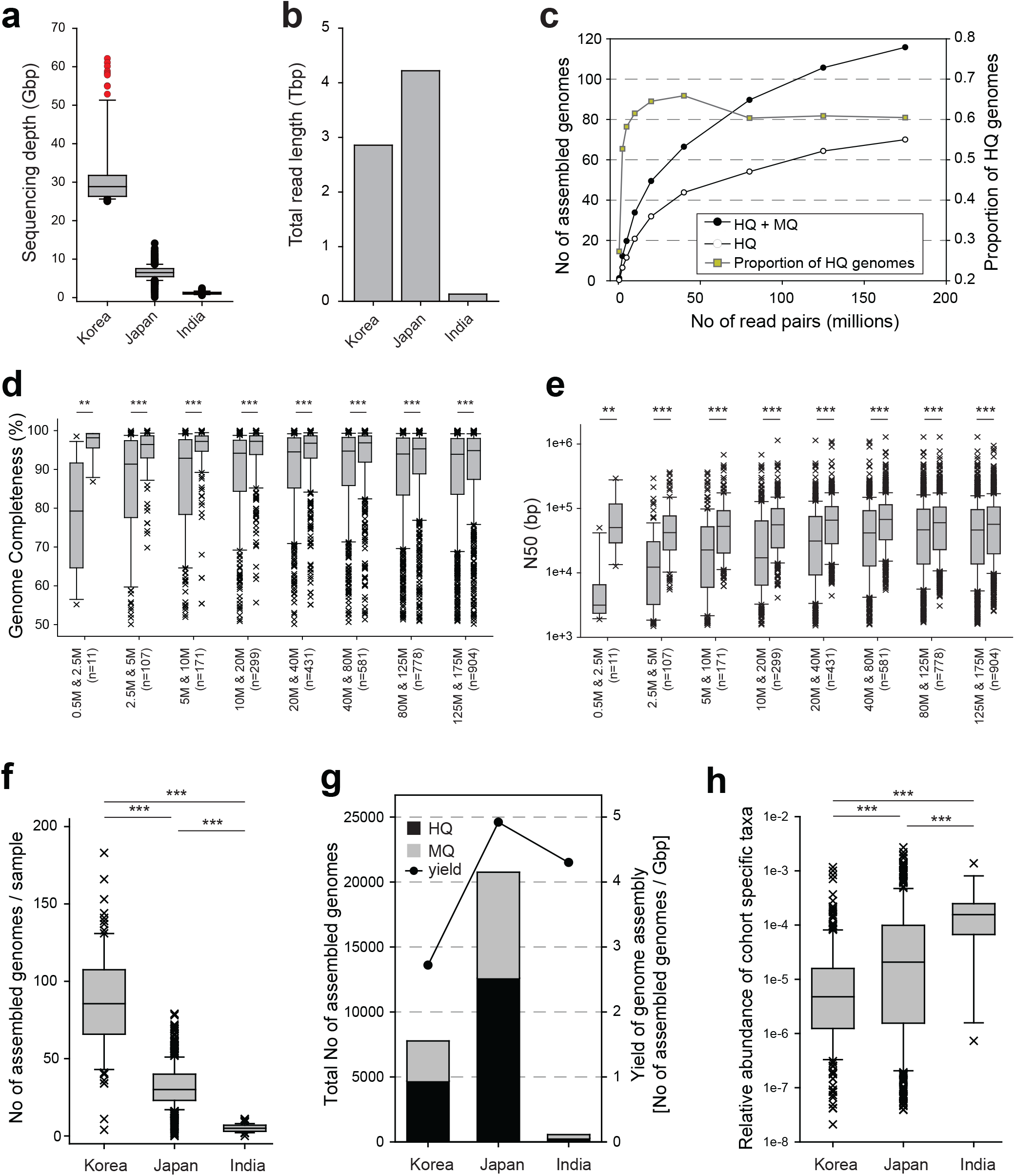
Effect of sequencing depth on *de novo* genome assembly. **a,** Sequencing depth of samples from Korea, Japan, and India. Red data points, nine samples used for the generation of simulated samples for different sequencing depths. **b,** Total read length of samples from Korea, Japan, and India. **c,** The average number of HQ and MQ genomes (left axis) and the proportion of HQ genomes (right axis) from nine samples. **d,e,** Completeness (d) and N50 (e) of assembled genomes from lower sequencing depth (left box of each column) and greater sequencing depth (right box of each column). **f,** The number of the assembled genomes from Korea, Japan, and India. **g,** Total number of the assembled genomes from Korea, Japan, and India, and genome assembly yields. **h,** The relative abundance of 224 Korea-specific, 338 Japan-specific, and 18 India-specific assembled genomes in independent fecal samples from the US (n = 926). *P*-values were calculated by two-sided Mann–Whitney U test (**: *P* < 0.01; ***: *P* < 0.001).

We then examined the effect of sequencing depth using the actual WMS data for KIJ samples. The number of HQ and MQ genomes assembled from each sample was highest in the ultra-deep sequenced samples from Korea (**Fig. 1f**). However, the proportion of HQ genomes in samples from Korea and Japan was not significantly different (**Fig. 1g; Supplementary Fig. 3c**). Notably, the genome assembly yield, i.e., the number of assembled genomes divided by the total sequencing length, was highest for samples from Japan (**Fig. 1g**). This suggests that sequencing hundreds of samples at a depth of 5–10 Gbp may constitute the most effective strategy for cataloging MAGs for a given population.

The ultra-deep sequencing may be advantageous for the genome assembly for low-abundance taxa. To test this, we compared MAGs exclusively assembled from each country but not included in the UHGG, i.e., 224, 388, and 18 genomes from Korea, Japan, and India, respectively. We then estimated their relative abundance in fecal samples in an independent population of 926 fecal samples from the US^16^, using Kraken2^17^. The genomes assembled exclusively from Korean samples shifted towards low-abundance taxa compared with genomes assembled from samples from other countries (**Fig. 1h**), which confirmed the original hypothesis.

### Cataloging reference genomes of 5,414 prokaryotic species from the human gut

To construct the most comprehensive reference database for the human gut microbiome, we integrated the newly generated 29,082 MAGs from KIJ samples with the UHGG genomes using dereplication approach (**Supplementary Fig. 1b**, **Methods**). Dereplication of the 29,082 MAGs resulted in 2,199 clusters of genomes. We selected a representative genome from each cluster to catalog the genomes for 2,199 representative species, which we then integrated with 4,644 representative genomes from the UHGG, via dereplication, resulting in 5,414 clusters of genomes. Finally, we selected 5,414 representative genomes and assigned their phylogenetic classifications using GTDB-Tk^18^ (**Fig. 2**). Among these representative genomes, 4,531 (83.7%) genomes were exclusively assembled from metagenomic data, which confirmed the notion that the major portion of the human gut microbiome has not yet been isolated. We identified 16S rRNA sequences in 2,542 representative genomes (47%) (**Supplementary Fig. 4**), covering the majority of phylogenetic clades. Unlike conventional databases of 16S rRNA sequences, the new database provides opportunities for functional interpretation of the detected taxa because it contains genomes corresponding to the 16S rRNA sequences.

**Fig. 2.**
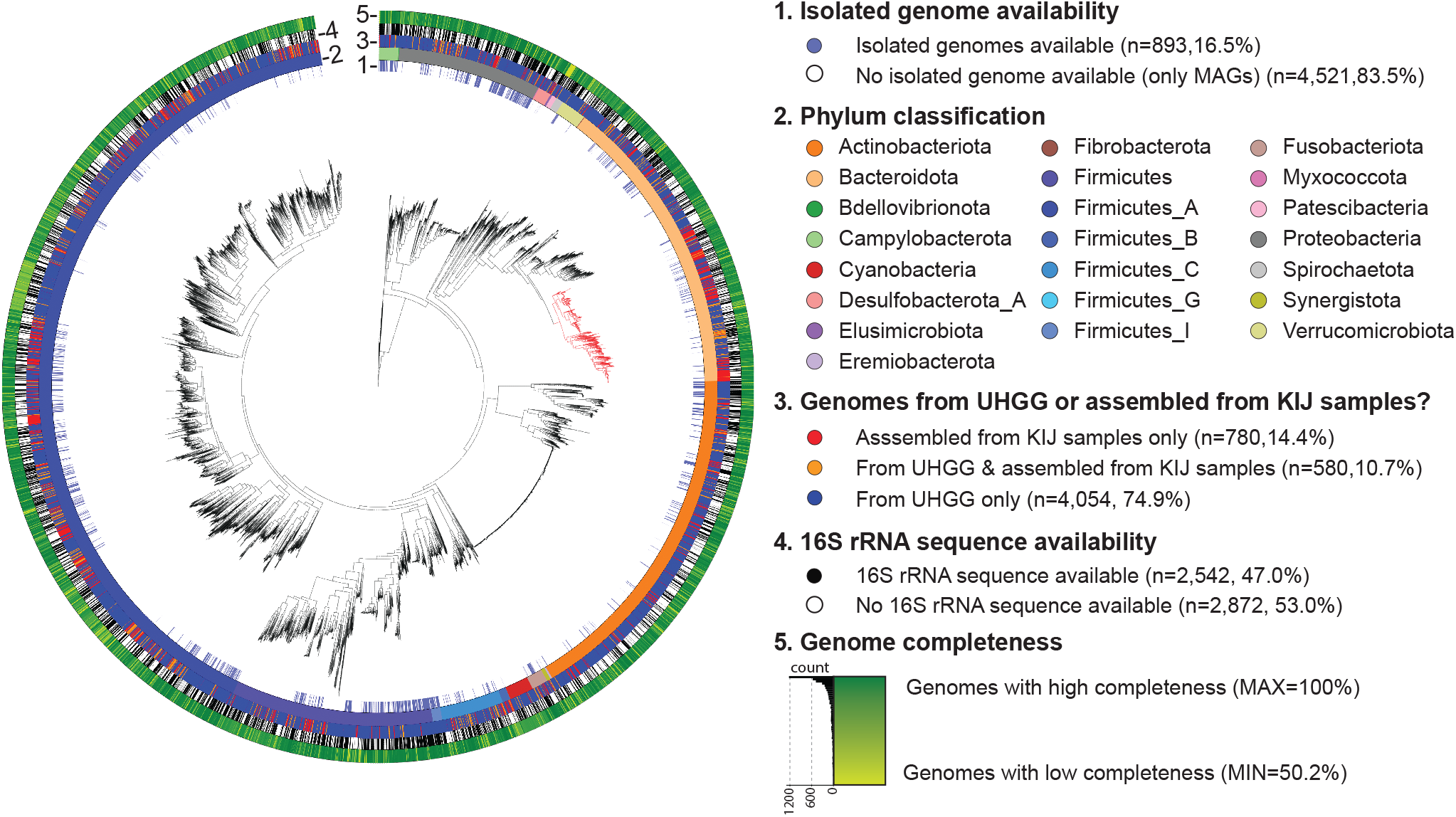
Phylogenetic tree of 5,386 representative genomes of prokaryotic species from the human gut contained in the HRGM. Maximum-likelihood phylogenetic tree reconstructed from 120 bacterial marker genes (**Methods**). Representative genomes were annotated by their isolated genome availability (1st layer from the inside), phylum classification (2nd layer), whether they were from UHGG or assembled from KIJ samples (3rd layer), 16S rRNA sequence availability (4th layer), and genome completeness (the outermost layer). Red branches represent 410 genomes from the *Bacteroidaceae* family that are enriched in the representative genome set updated by including KIJ samples.

The inclusion of MAGs from KIJ samples in the new database allowed several improvements on the UHGG. First, we reduced the data bias toward China among Asian countries (**Supplementary Fig. 5a**). Second, we expanded the total number of non-redundant reference genomes by 13.25% and the number of representative species by 16.6% increase (**Supplementary Table 3**). Among the 5,414 representative genomes, 780 genomes were assembled from KIJ samples only, and 536 representative genomes from the UHGG were replaced with new MAGs from KIJ samples. Hence, 1,316 representative genomes (28.3%) were updated in the HRGM (**Supplementary Fig. 5b**).

### New MAGs from Korea, India, and Japan are associated with diet-related lifestyles

Notably, *Bacteroidaceae* family (**Fig. 3,** redtree branches) was enriched in the updated MAGs (*P* < 0.001, Fisher’s exact test). Almost half the genomes from this family are from the *Bacteroides* genus and approximately two-thirds of the other half are from the *Prevotella* genus (**Supplementary Fig. 6**). Interestingly, three widely dispersed regions in the phylogenetic tree were highly enriched in the updated genome set. The first region (“a”) encompasses a portion of the *Prevotella* genus and includes 30 genomes annotated as *Prevotella copri*. Accordingly, westernized populations with a typically high-fat and low-complex carbohydrate diet exhibit low prevalence and diversity of *P. copri* compared with non-westernized populations^19^. The second region (“b”) encompasses a portion of the *Bacteroides* genus and includes 22 genomes annotated as *Bacteroides plebeius*. This species is typically found in Japanese subjects whose diet includes seaweed-rich food, such as sushi^20^. It has been suggested that *B. plebeius* harbors genes encoding an enzyme specific for algal carbohydrates, acquired from marine microbes. The third region (“c”) also encompasses a portion of the *Bacteroides* genus and includes 12 genomes annotated as *Bacteroides vulgatus*, which is typically present in the human distal gut, where undigested plant polysaccharides and proteins exist in large quantities^21^. Together, these observations indicate that the new MAGs from KIJ samples are associated with the diet-related lifestyles in Japan and Korea.

**Fig. 3.**
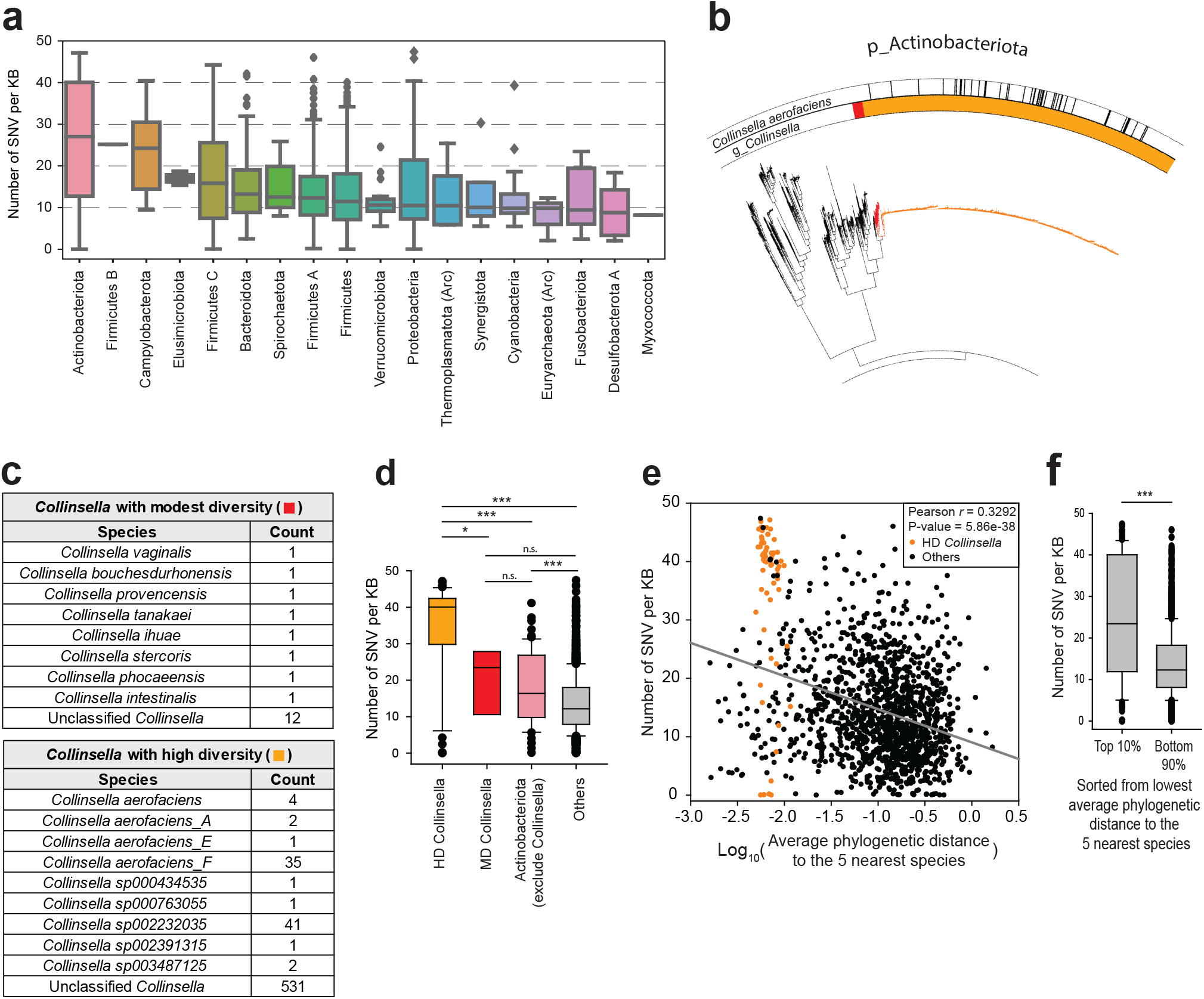
SNV density analysis of the relationship between within-species variation and speciation of gut microbes. **a,** The number of SNVs per kb pair of the aligned region. SNV density is summarized for each phylum. Boxes are sorted by the median. Arc, archaeal phylum. **b,** The phylogenetic tree for Actimobacteriota phylum. Inside annotation indicates the *Collinsella* genus, divided into *Collinsella* with modest phylogenetic dispersion (MD *Collinsella*, Red) and *Collinsella* with high phylogenetic dispersion (HD *Collinsella*, Orange). Black annotations in the outer circle represent *Collinsella aerofaciens*, *Collinsella aerofaciens_A*, *Collinsella aerofaciens_E*, and *Collinsella aerofaciens_F*, according to the GTDB-TK annotation. **c,** GTDB-TK based taxonomic annotation of MD *Collinsella* and HD *Collinsella*. **d,** SNV density of HD *Collinsella*, MD *Collinsella*, Non-*collinsella* actinobacteriota, and other species. **e,** Scatter plot analysis of SNV density and average phylogenetic distance to the five nearest species of each representative species. Orange points denote species of HD *Collinsella* and black points represent other species. **f,** Comparison of SNV density between the top 10% and bottom 90% species sorted from the lowest average phylogenetic distance to the five nearest species. Statistical significance was calculated by two-sided Mann–Whitney U test (n.s.: not significant; *: *P* < 0.05; ***: *P* < 0.001).

### SNV density is positively associated with the speciation rate of gut commensals

We then aligned genomes of species clusters containing ≥3 genomes with the representative genome and mapped SNVs (**Methods**). This yielded 274,543,071 SNVs from 2,821 species clusters, representing 10.07% and 13.34% increases, respectively, from the UHGG. The Actinobacteriota phylum had the highest SNV density (**Fig. 3a**). Phylogenetically overdispersed branches of Actinobacteriota species were apparent in both, the HRGM and UHGG. The majority of genomes from the overdispersed tree region belonged to the *Collinsella* genus. We divided these genomes into ones from a tree region with a modest phylogenetic dispersion (MD, 20 genomes) and those with a high phylogenetic dispersion (HD, 619 genomes) (**Fig. 3b**). Although the majority of genomes were not annotated at the species level, *Collinsella aerofaciens* was enriched in the HD group and other known *Collinsella* species were enriched in the MD group (**Fig. 3c**). SNV density in HD group was significantly higher than that of MD group (**Fig. 3d**).

SNV, a within-species genetic variation, is a major mechanism for the adaptation of commensal species to a distinct host environment. Wide dispersion of species branches indicates rapid speciation. Accordingly, high SNV density for a species with an overdispersed tree may indicate that the degree of within-species genetic variation may be positively associated with the speciation rate of gut commensals. To test this, we examined the correlation between SNV density of representative species and their phylogenetic distance to the five nearest species. The branch length to the neighboring species in the phylogenetic tree of a species that arose during rapid speciation tends to be short. We observed an inverse correlation between the average phylogenetic distance to the five nearest species and their SNV density (**Fig. 3e**), and a significantly higher SNV density for the top 10% species with shorter phylogenetic distance to the nearest five species than those for the bottom 90% species (**Fig. 3f**). This supports the model of a positive correlation of SNV density and the speciation rate of gut commensals.

### Functional landscape of 103 million proteins from human gut prokaryotes

Information on proteins encoded in the human gut microbes will facilitate the functional characterization of disease-associated microbiota. Using an in-house computational pipeline for cataloging human gut prokaryotic proteins (**Supplementary Fig. 1c** and **Supplementary Fig. 7**), we first identified 64,661,728 CDS (coding sequences) from 29,082 genomes from KIJ samples using Prodigal^22^. To reduce redundancy in the protein catalog, we first executed CD-HIT^23^ at 100% similarity level and then combined with proteins cataloged by the UHGP-100^11^. The consolidated protein catalog was next consecutively clustered by CD-HIT at lower sequence similarity levels: 95%, 90%, 70%, and 50%. This led to approximately 103.7, 20.0, 14.8, 8.5, and 4.7 million proteins at the sequence similarity levels of 100%, 95%, 90%, 70%, and 50%, respectively.

Unexpectedly, we observed that the UHGP contains proteins that are 100% identical, even in a catalog at 50% sequence similarity level. For instance, among the UHGP-50 proteins, GUT_GENOME232012_01109 and GUT_GENOME231777_00918 have an identical amino acid sequence. We identified 8,663, 82,507, 243,362, and 75,620,150 proteins that are redundant at 100% similarity in the UHGP-50, UHGP-90, UHGP-95, and UHGP-100, respectively. Exclusion of the UHGP proteins that were 100% identical revealed that the HRGM contains more proteins than UHGP at all levels of sequence similarity except for 50% (**Supplementary Table 3**).

To facilitate the functional interpretation of gut microbiome profiles, we next annotated functional genomic elements and proteins in the HRMG. We predicted and annotated non-coding RNAs and functional peptides, using Prokka^24^; antibiotic resistance genes, using RGI^25^; biosynthetic gene clusters, using antiSMASH^26^; and 16S rRNA regions, using barrnap^27^. For functional annotation of proteins, we used eggNOG-mapper^28^. Notably, the landscape of antibiotic resistance ontology revealed that phylogenetically close species in the human gut tend to share antibiotic resistance mechanisms (**Supplementary Fig. 8**). A significantly large portion of the human gut prokaryotic proteins has not yet been functionally annotated. For the HRGM protein catalogs at 100%, 95%, 90%, 70%, and 50% similarity levels, 13.13%, 28.05%, 29.17%, 36.35%, and 47.62% of proteins, respectively, had no functional annotation, according to eggNOG-mapper. This effect appears to be amplified by redundant proteins, resulting in a reduced annotation rate at low similarity level. Further, the annotation rate of proteins that are shared by many species is higher than that of species-specific proteins (**Supplementary Fig. 9**).

### HRGM improves taxonomic and functional classification of sequencing reads

According to a recent benchmark study, whole-DNA–based methods outperform marker-based methods for taxonomic classification of metagenomic sequencing reads^29^. The performance of whole-DNA–based methods relies on the quality of the reference genome database. The standard databases lack numerous genomes of species that exist in the human gut, which leads to false-negatives, while including many genomes from other microbial communities, which leads to false-positives^29^. We hypothesized that the HRGM, which is specific to the human gut microbiome and more comprehensive than other databases, can improve the taxonomic classification of sequencing reads. We used Kraken2^17^ to compare the taxonomic classification of three genome databases: a standard database that contains RefSeq^30^ complete genomes (RefSeq CG) of bacterial, archaeal, and viral domains; the UHGG-based database; and the HRGM-based database. To generate independent test datasets, we compiled WMS data for 1,022 fecal samples from the US, Cameroon, Luxembourg, and Korea, which were not included in the UHGG nor HRGM. We then evaluated the efficacy of Kraken2 classification based on the proportion of classified reads (**Methods**). The classification efficacy using the UHGG and HRGM-based databases was substantially higher than that of the standard database (**Fig. 4a,b**, *P* < 0.001, two-sided Wilcoxon signed-rank test). In addition, the variance of the read classification rate of custom databases was significantly smaller than that of the standard database, except for the Cameroon population (**Fig. 4a**, *P* < 0.001, Brown-Forsythe test). Importantly, the classification efficacy of the HRGM-based database was significantly improved compared with that of the UHGG-based database for the four test samples (**Fig. 4a,c**, *P* < 0.001, two-sided Wilcoxon signed-rank test), which suggests that the updated reference genome database improves taxonomic classification of the gut metagenomic sequencing data.

**Fig. 4.**
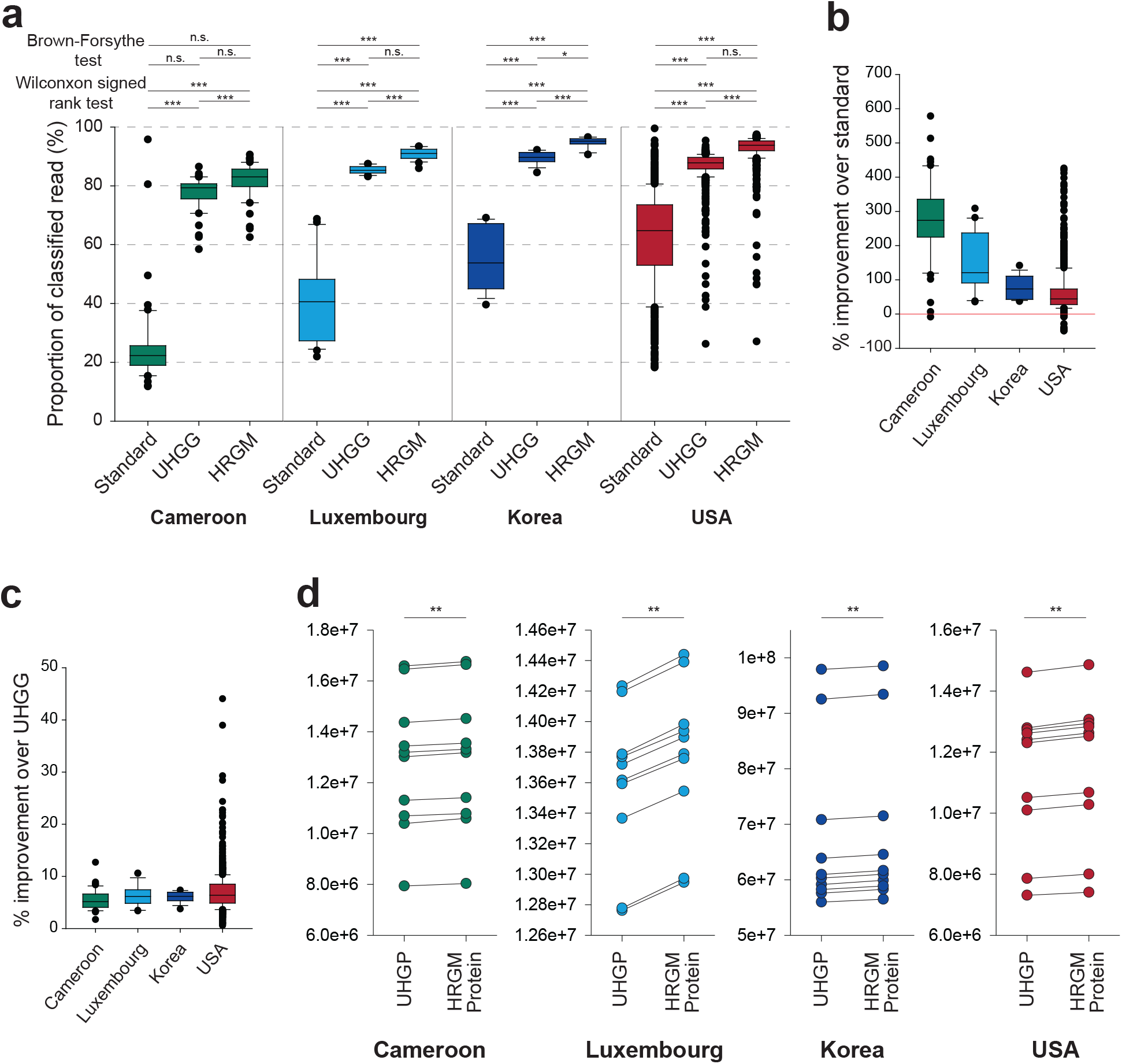
Effect of HRGM on taxonomic and functional classification of sequencing reads. **a,** Proportion of taxonomically classified sequencing reads of WMS data from four different populations. The significance of the improvement was calculated by Wilcoxon signed-rank test. Brown–Forsythe test was used to evaluate the decrease of variance. **b,c,** Percent improvement of the read classification proportion in HRGM-based database compared with the standard database (b) and the UHGG-based database (c). **d,** The number of reads aligned to the UHGP-95 and HRGM-95 protein catalogs. Statistical significance was calculated by using Wilcoxon signed-rank test.

Next, we investigated the efficacy of functional classification based on the number of aligned sequencing reads from reference protein databases. Because of the extremely large number of reference proteins, we used only 40 samples randomly selected from the 1,022 fecal samples (10 samples from each population), and aligned the sequencing reads with the UHGP-95 and HRGM-95 protein catalogs (**Methods**). The number of aligned reads was 1.31% higher, on average, with HRGM-95 in all tested samples than with UHGP-95 (**Fig. 4d**), although HRGM-95 contains 0.4% more proteins than UHGP-95.

Taken together, the newly assembled genomes from under-represented Asian countries significantly improve the genome and protein databases for metagenomic analysis of both, taxonomic and functional profiling.

### Reliable taxonomic profiling of low-abundance taxa requires deep sequencing

Taxonomic profiles obtained by shallow sequencing (0.5–2 million reads) highly correlate with those obtained by ultra-deep sequencing (2.5 billion reads)^31^. However, this evaluation is based on entire taxa, in which highly abundant or core taxa govern the correlation measure. Further, low-abundance taxa likely play important, as yet unknown, biological roles in the gut microbial communities^32,33^. We therefore evaluated the impact of sequencing depth on the reliability of taxonomic profiling for different ranges of taxon abundance. We generated a simulated dataset at various sequencing depths 16 new Korean fecal samples, and not included in the HRGM. We then stratified the taxonomic features into eight different groups, according to the mean relative abundance (**Fig. 5a,b**). We calculated the mean Pearson correlation coefficient (*PCC*) and the mean Spearman correlation coefficient (*SCC*) between the taxonomic profiles at different sequencing depths for different mean relative abundances (**Methods**). The taxonomic profile similarity between two groups showed increasing *PCC* and *SCC* with an increasing sequencing depth. For example, >10 million read pairs (3 Gbp) may need to have taxonomic profiles that highly correlate (*PCC* > 0.9) with those based on 80 million read pairs (25 Gbp) to account for the features with lowest 13.92% of relative abundance (relative abundance < 1e–06) (**Fig. 5c and Supplementary Fig. 10a**). For *SCC* > 0.9, the required sequencing depth increased to 20 million read pairs (6 Gbp) for taxonomic features with a similar level of relative abundance (**Fig. 5b and Supplementary Fig. 10b**). Overall, these observations suggest that deep sequencing (>20 million read pairs) may be required to obtain reliable taxonomic profiles of low-abundance taxa.

**Fig. 5.**
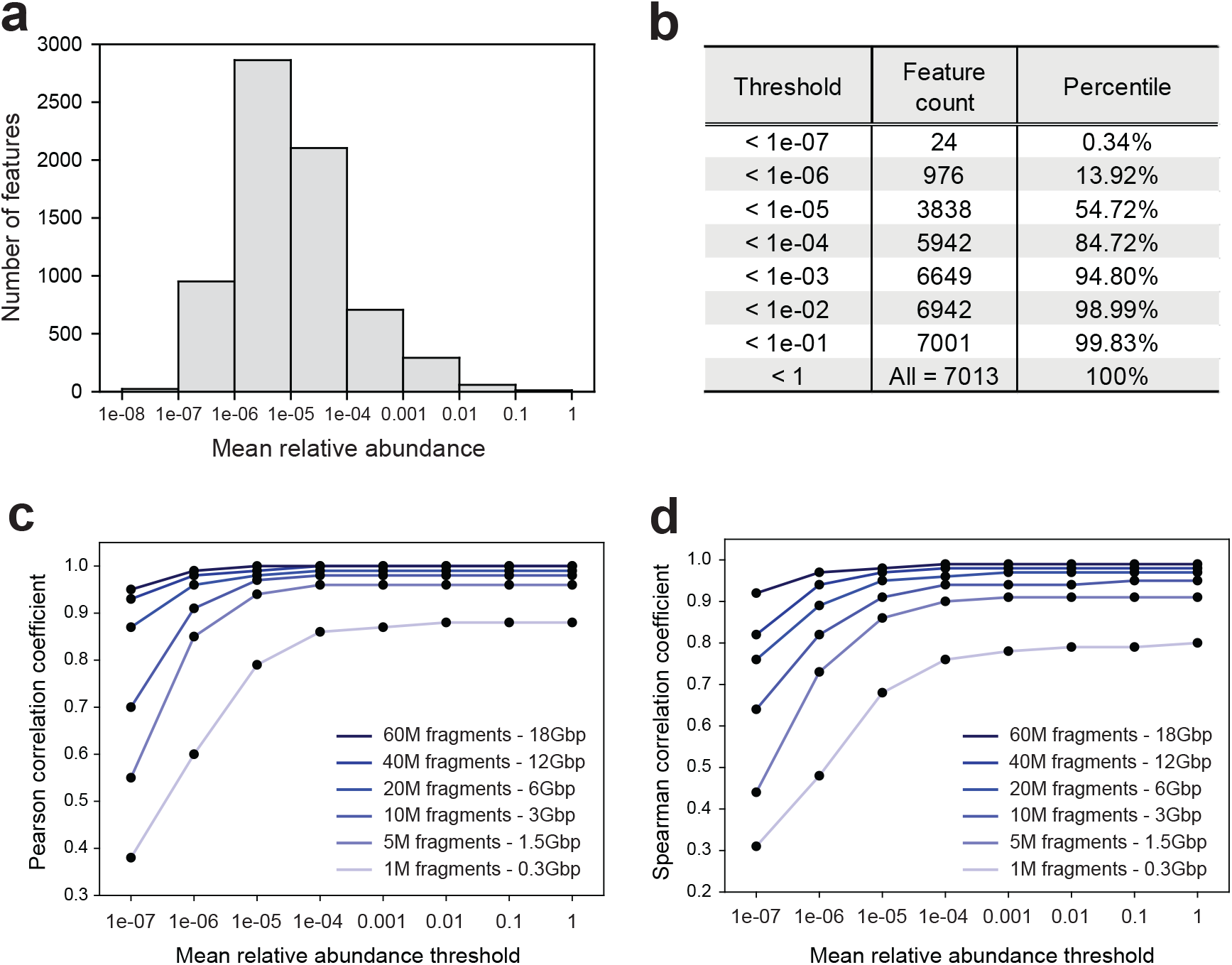
Effect of sequencing depth on the reliability of taxonomic profiles. **a**, The distribution of taxonomic features over different mean relative abundances. **b**, The cumulative proportion of taxonomic features at different thresholds of mean relative abundance. **c,d,** Pearson correlation coefficient (*PCC*) (c) and Spearman correlation coefficient (*SCC*) (d) of the taxonomic profiles at the given sequencing depth and 80M fragments. The x-axis (the mean relative abundance threshold) indicates the upper boundary of the mean relative abundance.

### Sequencing 30 Gbp is optimal for functional profiling of the human gut microbiome

Next, using the protein catalog, we investigated the optimal sequencing depth for functional profiling of the human gut microbiome. Since the detection of gene content generally requires a much deeper sequencing depth than that for the detection of genomes, we analyzed the WMS data for five Korean fecal samples at a depth of approximately 200 million read pairs (60 Gbp) (**Methods**). The number of the detected coding genes initially grew rapidly as the sequencing depth increased, but later approached the estimated maximum count (**Supplementary Fig. 11a**). The curves fitted well (*R*^2^ > 0.99) two-site saturation models^34^, and we hence estimated the maximum number of coding genes for each sample using the regression model. Interestingly, the estimated maximum gene counts in the samples differed, reflecting the different alpha diversity of the microbial community. However, all samples showed very similar normalized maximum gene count curves, with over 80% of the gut microbial coding genes detected by sequencing 30 Gbp or 100 million read pairs in all samples (**Supplementary Fig. 11b**). Sequencing another 30 Gbp would fail to detect 90% of the maximum gene count. Therefore, 100 million read pairs is the optimal sequencing depth for the best trade-off between the sequencing cost and the gain-of-functional information for WMS-based studies of the human gut microbiome.

### Profiling cross-reactivity potential identifies autoimmune-associated commensals

Microbial peptides homologous to the host auto-antigens may stimulate host immune cells and, hence, the hypothesis of molecular mimicry has emerged as a mechanism underlying autoimmune diseases^35^. To systematically evaluate this hypothesis, we mapped microbial peptide sequences homologous to the human self-antigens involved in autoimmune diseases onto the genomes of HRGM representative species. We first compiled autoimmune disease-related antigen set from the Immune Epitope Database (IEDB)^36^, and then used it for homology-searches of microbial peptide sequences from 5,414 representative species. We thus identified species with a high cross-reactivity potential based on the density of the encoded cross-reactive epitopes. Because the number of epitope-containing genes (ECG) increased as the number of coding genes increased (**Fig. 6a**), we divided the ECG count by the total number of genes for each species. Some human gut commensals had a relatively high cross-reactivity potential (**Fig. 6b,c, Methods**). On the genus level, *Akkermansia*, *Alistipes*, *Bifidobacterium*, *Lawsonibacter*, *Oscillibacter*, *Prevotella*, and *Sutterella* have a high cross-reactivity potential (**Fig. 6d**). Indeed, many of them are associated with autoimmune diseases. For example, *Akkermansia muciniphila* is abundant in the enthesitis-related arthritis patients^37^, while *Bifidobacterium* is enriched in these^37^ and inflammatory bowel disease (IBD) patients^38^. Increased abundance of *Oscillibacter* is accompanied by increased levels of interleukin 6^39^, a pro-inflammatory cytokine that can disrupt the immune homeostasis and increase the risk of autoimmune diseases. The abundance of intestinal *Prevotella copri* is strongly correlated with the risk of arthritis^40^ and *Sutterella wadsworthensis* is enriched in ulcerative colitis patients who do not respond to fecal microbiota transplantation^41^. These suggests that cross-reactivity potential of commensal genomes is predictive for human gut microbiota associated with autoimmune diseases.

**Fig. 6.**
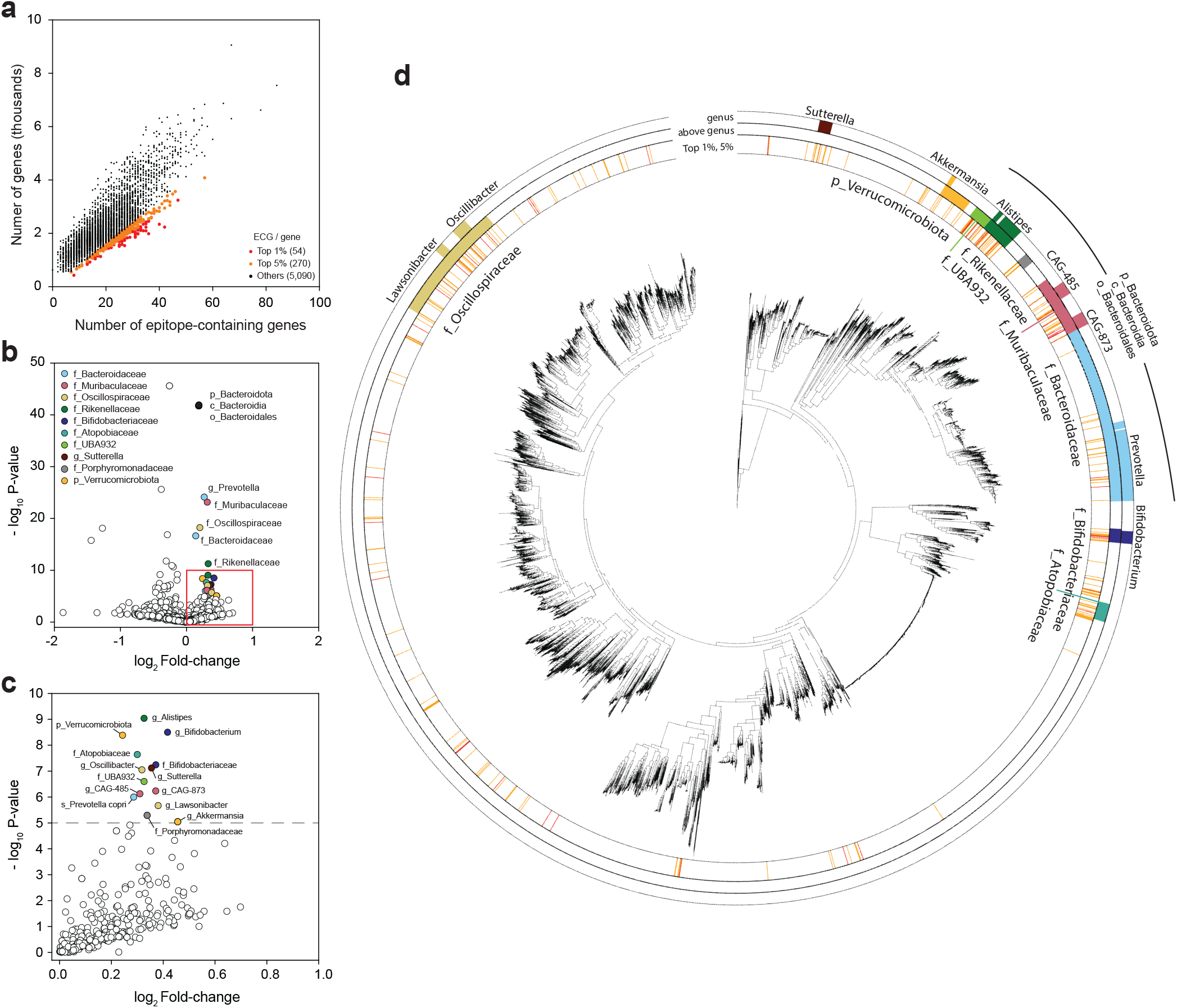
Landscape of cross-reactivity potential of gut prokaryotic genomes. **a,** The number of genes and autoimmune epitope sequence-containing genes (ECG) in 5,414 genomes of species representatives. Red and orange points, species with the top 1% and 5% ECG per gene, respectively. **b,** Volcano plot of the enrichment of ECG density. Taxonomic clades with positive log2 fold-change and *P* < 1e–5 are highlighted with different colors. Taxonomic clades denoted by the same color have an inclusive relationship (e.g., *g_Prevotella* belongs to *f_Bacteroidaceae*), with the exception of p_Bacteroidota, c_Bacteroidia, and o_Bacteroidales. The first character of each clade name indicates the taxonomic levels (p: phylum; c: class; o: order; f: family; and g: genus). **c,** The red-highlighted area from (b). **d,** Maximum-likelihood phylogenetic tree with taxonomic annotations of clades with high ECG density. The first layer represents clades with the top 1% (red) and 5% (orange) ECG density [annotations and color designations are the same as in (**a**)]. The second and third layers represent enriched taxonomic clades in the volcano plot [taxonomic annotations and color designations are the same as in (b) and (c)]. The second layer represents above-genus level annotations. The third layer represents genus-level taxonomic clades.

## Discussion

In the present study, we constructed an improved catalog of the human reference gut prokaryotic genomes and their proteins, by including MAGs from fecal metagenomes from under-represented Asian countries. Inclusion of the newly assembled genomes expanded the catalog size by over 10%. In addition, we demonstrated that database expansion also significantly improved the taxonomic and functional classification of sequencing reads. Many new MAGs were associated with diet-related lifestyles at the sampled geographic locations.

Therefore, complementation of metagenome datasets to account for under-sampled geographical locations and lifestyles might be an effective strategy for improving the human reference gut microbiome.

We also demonstrated that the analysis of microbial DNA and peptide sequences facilitates the understanding of gut commensal speciation and interactions with the host immunity. The colonizing commensal microbes adjust to their host environment via genetic changes and selection, which lead to genetic variation within species. We cataloged the SNVs of conspecific genomes and found that the SNV density of gut prokaryotic species is inversely correlated with the phylogenetic distance to their neighboring species. This may suggest that the degree of within-species genetic variation is positively associated with the speciation rate of gut commensal microbes. Whether SNV actually enhances the speciation rate should be addressed in future investigations. Finally, we showed that systematic analysis of microbial peptide sequences homologous to the host auto-antigens allows the prediction of gut microbial taxa potentially associated with autoimmune disease via the mechanism of molecular mimicry. Such analysis is only possible if microbial protein sequences are available with the corresponding taxonomic information.

As the WMS analysis for population-wide human gut microbiome profiling increases in popularity, the choice of sequencing depth is an important factor to consider in study design. Here, we demonstrated that deep sequencing (>20 million read pairs) is necessary for reliable taxonomic profiling of low-abundance commensals. The current knowledge of human gut microbiome is biased towards core taxa that are usually highly abundant. Low sequencing depth (e.g., 0.5–2 million read pairs) may be sufficient for the profiling of core taxa, but not those with low abundance. Deep sequencing may therefore be required for the WMS-based analysis of human gut microbiome to investigate the function of relatively unexplored low-abundance species. Accordingly, the current study provides the guidelines for the choice of sequencing-depth for the analysis of human gut microbiome for different purposes.

In conclusion, the HRGM database, which contains information on various biological entities, from DNA and protein sequences to pan-genomes of species, is a versatile resource for functional dissection of disease-associated gut microbiota. The data will be available via a web server (www.mbiomenet.org/HRGM/) and will be periodically updated as new WMS data for fecal samples become publically available.

## Methods

### Sequencing fecal metagenome samples from Korea, India, and Japan

WMS data for fecal samples from India and Japan were obtained from published studies^14,15^. Fecal WMS data for India were generated for 110 healthy donors in North-Central and Southern India^14^. Although the sequencing depth was relatively low (1.2 Gbp on average), it was expected that many novel genomes would be assembled because MAGs from India are not included in the existing catalogs. By contrast, 805 MAGs from Japan are included in the UHGG. However, it was expected that the inclusion of the recently published deep-sequencing WMS data for 645 Japanese fecal samples (6.5 Gbp on average)^15^ would greatly expand the number of MAGs for Japan. In addition, ultra-deep WMS data (31 Gbp on average) were generated for fecal samples from 90 Koreans recruited by the Severance Hospital (Seoul, Korea; IRB No 4-2020-0309 and IRB No 4-2017-0788). Written informed consent was obtained before the study. The UHGG does not contain any MAGs from Korea.

The libraries were prepared as described in the TruSeq Nano DNA Library Prep Reference Guide (Illumina #15041110). Briefly, 100 ng DNA was fragmented using LE220 Focused ultrasonicator (Covaris, Inc.). Fragmented DNA was end-repaired and approximately 350-bp fragments were obtained after size selection. After adapter ligation, eight PCR cycles were performed. Library quantification was performed as described in the Kapa Illumina Library Quantification Kit (Kapa Biosystems, #KK4854). Next, 150 bp ×2 paired-end sequencing was performed using Illumina HiSeq4000. In summary, new WMS data for 845 fecal samples collected from Korea, India, and Japan were obtained. The total read length was 7.2 Tbp. All samples used in the current study are described in **Supplementary Table 2**.

### Metagenome assembly and binning

The adapter sequences were trimmed, and low-quality bases and short reads were removed from WMS data using Trimmomatic v0.39^42^. Next, the reads were aligned with the human genome GRCh38.p7 using Bowtie2 v2.3.5^43^, and the aligned reads were then removed. The majority of quality-controlled reads were assembled as contigs using metaSPAdes^44^, which is a metagenome-specific pipeline of SPAdes v3.13.0. For unknown reasons, and regardless of sample size, metaSPAdes runtime was excessively long for 107 samples. In those cases, MEGAHIT v1.2.8^45^ was used (**Supplementary Table 2**).

Genome bins were generated using the ensemble approach and three binning tools: MetaBAT2 v2.13^46^, MaxBin2.0 v2.2.6^47^, and CONCOCT v1.1.0^48^. First, the reads from each sample were first aligned with the assembled contigs from the previous step using Bowtie2, and the three binning programs were initiated. The minimum size of a contig for binning was set at 1,000 bp, except for MetaBAT2, which requires at least 1,500 bp. The three binning predictions were combined for improved binning results using the bin refinement module of MetaWRAP v1.2.2^12^, which uses CheckM v1.0.18^49^ to evaluate the quality of genome bins in terms of completeness and contamination rate. The minimum completeness was set at 50%, the maximum contamination at 5%, and the minimum quality score (*Completeness* - *5* × *Contamination*) at 50. The same threshold values for CheckM results were applied during the construction of the UHGG. This resulted in 7,767 genomes from Korean samples, 563 genomes from Indian samples, and 20,752 genomes from Japanese samples (29,082 genomes in total). The genome bins were divided into two groups: HQ, bins with over 90% completeness and less than 5% contamination; and MQ, bins with 50–90% completeness and less than 5% contamination.

### Generation of genomic species clusters

Groups of genomes that corresponded to species were generated using a two-step iterative procedure. Preliminary clustering was performed using Mash v2.2^50^ algorithm. Mash distances were calculated for all possible pairs of genomes using the “-s 10,000” parameter. Next, the average-linkage–based hierarchical clustering was performed, at a cutoff of 0.2. Mash algorithm is sufficiently fast to calculate all-by-all distances for hundreds of thousands of genomes in a timely manner. However, this compromises the accuracy, especially for low-coverage genome pairs^51^, which are common in MAGs. Therefore, to improve cluster quality, ANImf^51^ was calculated for every pair of genomes within each initial cluster. To avoid the over-estimation of ANI by local alignment, a minimum coverage threshold was applied for each pair. The coverage cutoff of genome A and genome B was determined at *min(0.8, Completeness of genome A* × *Completeness of genome B)*. If the alignment coverage between two genomes was lower than the cutoff, they were regarded as different genomes. The genomes were then clustered using the average linkage-based hierarchical clustering at a cutoff of 0.05 (or 95% identity), which is a widely accepted ANI threshold for species-level boundary^4,9–11,52^. The genome intactness score (*S*)^9,11^, *S* = *Completeness* − *5* × *Contamination* + *0.5* × *log*_*10*_*(N50)*, was then calculated. For clusters containing more than two genomes, a genome with the highest *S* was selected as the representative genome for the cluster. The above two-step procedure was iterated until the clusters ceased to change. Hence, 2,199 species clusters were generated for 29,082 genomes from KIJ samples, with eight iterations of the aforementioned procedure (**Supplementary Fig. 1a**). Finally, the 2,199 genomes were combined with 4,644 genomes from the UHGG, generating 5,414 species clusters for the HRGM at the fourth iteration (**Supplementary Fig. 1b**).

### Non-redundant genome counting

To count the number of non-redundant genomes, the redundant genomes were removed, similar to what was done for the UHGG pipeline^11^. First, the pairwise genome distance was calculated using Mash^50^ and the entire genomes were clustered using average-linkage–based hierarchical clustering, with a 0.001 cutoff (Mash ANI 99.9%). To reduce the computation time, the hierarchical clustering was performed only for the connected components with the distance of 0.1, because it is highly unlikely that genomes that are not within the distance of 0.1 are clustered together by a distance of 0.001. In the process, 22,761 genomes were clustered into 8,508 conspecific genome bins. Multiple genomes from the same sample in the same species bin were counted only once.

### Taxonomic and functional annotation of representative species genomes

The taxonomic annotation of 5,414 representative species genomes was performed using the “classify_wf” function of GTDB-Tk v1.0.2^18^. The reference version was GTDB R04-RS89, released in June 2019. Genomic features, such as CDS, rRNA, and tRNA, were identified and annotated in each genome using Prokka v1.14.5^24^ with “--kingdom Bacteria” and “--kingdom Archaea” parameters for the bacterial and archaeal genomes, respectively. With the protein sequences predicted by Prokka, the antibiotic resistance genes were annotated using RGI v5.1.0^25^ with default parameters. The landscape of antibiotic resistance potential of 5,414 species-representative genomes is depicted in **Supplementary Fig. 8**. Finally, the secondary metabolite gene cluster was annotated using antiSMASH v5.1.2^26^. For the full-featured annotation, the “--cb-general, --cb-knownclusters, --cb-subclusters, --asf, --pfam2go, --smcog-trees, --cf-create-clusters” parameters were set.

To render the HRGM useful for the 16S rRNA sequencing-based metagenomic analysis, the 16S rRNA regions for 5,414 representative species genomes were predicted using barrnap v0.9^27^ tool and the “--evalue 1e-05” parameter, and “--kingdom bac” and “--kingdom arc” parameters for bacterial and archaeal genomes, respectively. The 16S rRNA sequences were thus directly predicted from 1,364 representative species genomes. For the remaining 4,050 representative species, the search for 16S rRNA sequences was expanded to their conspecific genomes. The barrnap analysis was used for the genomes from KIJ samples and pre-established 16S rRNA region annotations were used for the genomes from the UHGG. Within the expanded search space, 16S rRNA sequences were identified for 1,178 additional genomes. Consequently, 16S rRNA sequences were generated for 2,542 species in the HRGM (**Supplementary Fig. 4**).

### Cataloging SNVs

For the species bins with more than three genomes, SNVs were identified using the codes provided by the UHGG^11^. Briefly, non-representative genomes were aligned with the representative genome in the species bin using nucmer 4.0.0beta2^53^. Best bi-directional alignments were identified using the delta-filter program and “-q –r” options, and SNVs were annotated using the show-snp program; nucmer, delta-filter, and show-snp are software packages of MUMmer v3^54^. For each species bin (*G*) whose representative genome is *r*, the number of SNV per kb was calculated as follows:

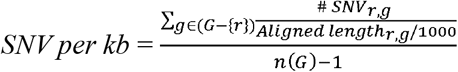

*SNV per kb* was only calculated for 1,521 species bins with ≥10 genomes to avoid bias. For the 1,521 genomes, the average phylogenetic distance to the five nearest species was calculated using the IQ-Tree^55^.

### Cataloging gut prokaryotic proteins and their functional annotation

Overall, 64,661,728 CDS were identified in 29,082 genomes from the KIJ set using Prodigal v2.6.3^22^ and “-c -m -p single” parameters. Since many proteins were derived from conspecific genomes, the catalog may have included many homologous proteins. To reduce the redundancy in the protein catalog, CD-HIT v4.8.1^23^ was adopted. To reduce CD-HIT running time, identical proteins were first clustered and then CD-HIT was executed at 100% similarity level. The cataloged proteins were then combined with those in UHGP-100^11^. The consolidated protein catalog was subsequently submitted to CD-HIT clustering analysis at five different sequence similarity levels, 100%, 95%, 90%, 70%, and 50%. For accurate and efficient clustering, a multi-step iterative clustering method recommended by the CD-HIT tutorial was adopted. For instance, the CD-HIT-95 protein catalog (a 95% similarity level protein catalog) was constructed based on CD-HIT-100 proteins, and the CD-HIT-90 protein catalog was constructed based on CD-HIT-95 proteins. This resulted in approximately 103.7 million, 20.0 million, 14.8 million, 8.5 million, and 4.7 million proteins at the sequence similarity levels of 100%, 95%, 90%, 70%, and 50%, respectively. The overall pipeline for protein catalog construction is depicted in **Supplementary Fig. 7**.

Representative protein sequences in the five protein catalogs were functionally annotated using eggNOG-mapper v2.0.1^28^, which is based on the eggNOG protein database v5.0^56^. The resultant annotations include eggNOG orthologs and functional terms from several databases, including Gene Ontology (GO)^57^ and Kyoto Encyclopedia of Genes and Genomes (KEGG)^58^. Further, for each protein cluster, taxonomic origins of all member proteins and the lowest common ancestor of the cluster were tracked and annotated.

The numbers of shared species and shared phyla of proteins in the HRGM-50 protein catalogs were annotated based on the taxonomic annotation of member proteins. The number of shared species was binned at the bin size of 10, then the annotation rate for each protein bin was calculated as the number of annotated proteins divided by the number of proteins in the bin.

### Reconstruction of the phylogenetic tree

For the bacterial and archaeal genomes, 120 and 122 universal marker genes, respectively, were predicted by the GTDB-Tk^18^. Using the concatenated sequences of marker genes, the maximum-likelihood tree was generated using IQ-TREE^55^. The phylogenetic tree of bacterial genomes was visualized using iTOL^59^.

### Kraken2 databases

The Kraken2 v2.0.8-beta^17^ custom database for the HRGM representative genomes was prepared based on the taxonomic annotations in GTDB-TK^18^. When two or more genomes were annotated to the same taxon, they were discriminated at the succeeding lower rank. For example, if *genome a* and *genome b* were both annotated to *species_A*, *genome a* and *genome b* were annotated as *Species_A;strain_1* and *Species_A;strain_2*, respectively. By doing so, the user can select a taxonomic rank, thereby measuring species abundances together or individually.

The Kraken2 database for the UHGG^11^ was downloaded from UHGG FTP on March 6, 2020. The Kraken2 standard database was downloaded and constructed using “kraken2-build --standard” command on July 14, 2020.

### Measuring taxonomic classification rate of sequencing reads

WMS data were compiled for publicly available data for 926, 54, and 26 fecal samples from the US^16^, Cameroon^60^, and Luxembourg^61,62^, respectively. WMS data for 16 fecal samples collected from Korea, which were not included in the HRGM, were also used. These 1,022 fecal samples were neither used for the UHGG nor for the HRGM. The data were pre-processed and taxonomically classified using Kraken2 with standard database, UHGG-based database, and HRGM-based database. The taxonomic classification rate was then calculated based on the proportion of aligned sequence reads in a sample with respect to the database.

### Measurement of functional classification rate of sequencing reads

The functional classification rate of sequencing reads was determined based on the number of aligned reads against the protein catalog. For the analysis, WMS data were randomly selected for ten fecal samples from the Cameroon, Korea, US, and Luxembourg cohorts (the same samples were used for the taxonomic classification assessment). After pre-processing, 40 samples were aligned with the UHGP-95 and HRGM-95 protein databases using blastx of DIAMOND v0.9.35.136 ^63^. The results were filtered at >80% query coverage (read coverage) and >95% alignment identity thresholds. A pair of reads was treated as two independent reads. For multiple alignments of a read, only the best alignments by bit score and e-value were considered.

### Finding the optimal sequencing depth for gene-level analysis of the gut microbiome

For five Korean fecal samples, WMS data generated at a sequencing depth of >60 Gbp, the reads were aligned against the HRGM-95 protein database using blastx of DIAMOND^63^. Alignment results with >80% read coverage and 80% identity were included in further analysis. For each sample, the number of detected genes with at least one aligned read was counted by iteratively removing 1000 randomly selected reads. The number of the detected genes for a given sequencing depth exhibited a saturation curve. The curve fitted well (*R*^2^ > 0.99 for all samples) the two-site binding model^34^. The required sequencing depth for a given gene coverage was determined based on the estimated maximum number of genes according to the equation.

### Evaluation of the effect of sequencing depth on *de novo* genome assembly

Nine Korean samples with sequencing depth of >52.5 Gbp (**Supplementary Table 2**) were selected for analysis. Then, 0.5, 2.5, 5, 10, 20, 40, 80, 125, and 175 million read pairs were randomly sampled from each of these samples. As the average read-pair length was 300 bp, the sequencing depths of these random samples corresponded to 150 Mbp, 750 Mbp, 1.5 Gbp, 3 Gbp, 6 Gbp, 12 Gbp, 24 Gbp, 37.5 Gbp, and 52.5 Gbp, respectively (**Supplementary Fig. 2**). For the 81 simulated samples (9 samples × 9 depths), *de novo* genome assembly was performed using the same pipeline as that used for the database construction.

Two adjacent sequencing depths (e.g., 125 *vs.* 175 million read pairs) were compared to evaluate the effect of sequencing depth on the *de novo* genome assembly. Samples with a greater sequencing depth may yield more MAGs with over 50% completeness, yet with a lower average quality, than those with a lower sequencing depth because of MAGs that barely pass the completeness threshold. Therefore, instead of the average quality scores of all assembled genomes, two genomes assembled at different sequencing depths for the same species clusters were compared. Mash^50^ clustering of genomes from two random samples was performed for a comparison based on the average-linkage–based hierarchical clustering, at a threshold of 0.1 (90% identity). Mash clustering was sufficient for clustering conspecific genomes in the simulated samples. Indeed, no cluster had more than two genomes from the same sequencing depth. The assembly quality (completeness, contamination, N50, and genome size) of conspecific genomes at adjacent sequencing depths was then compared.

### Evaluation of the effect of sequencing depth on taxonomic profiling

To avoid overestimation of performance, WMS data for 16 Korean fecal samples that have not been used for the HRGM construction and generated at a sequencing depth of >24.5 Gbp were used. From each of the 16 samples, 1, 5, 10, 20, 40, 60, and 80 million read pairs that corresponded to 300 Mbp, 1.5 Gbp, 3 Gbp, 6 Gbp, 12 Gbp, 18 Gbp and 24 Gbp, respectively, were randomly sampled. Taxonomic profiling was then conducted using Kraken2 and the HRGM-based database. Based on the hypothesis that profiling of low-abundance taxa is more affected by sequencing depth than abundant ones, the taxonomic features were stratified at eight different levels of relative abundance, ranging from 1e–07 to 1 with every ten-fold increase (**Fig. 5a,b**). *PCC* and *SCC* between the taxonomic profiles at different sequencing depths were then calculated for each group of features for different levels of relative abundance.

### Profiling cross-reactivity potential of the gut prokaryotic genomes

Epitope sequences from autoimmune disease-related self-antigen were compiled from IEDB^36^. “Epitope: Linear epitope”, “Antigen: Organism: Homo sapiens”, “Host: Homo sapiens”, and “Disease: Autoimmune Disease” filters from the IEDB web portal were applied. Epitope sequences that required post-translational modification (e.g., citrullination and deamination) and epitopes shorter than five amino acids were removed. Next, 24,461 unique epitope sequences were aligned with the protein sequences encoded by 5,414 species representatives using BLASTP ^64^. For meticulous alignment of short peptide sequences, “-word_size 4”, “-evalue 10000”, and “-max_target_seqs 100000” options were applied. For every epitope-to-gene pairwise alignment, the Alignment Score (*AS*) was calculated, as follows:

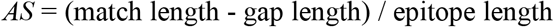

*AS* = 1 alignments were used and the number of protein-coding genes of autoimmune disease epitopes was calculated for every representative species. The number of ECGs was positively correlated with the number of genes. Therefore, the number of ECGs was normalized to the number of genes. To identify epitope-enriched taxonomic clades, EGC per gene of each taxonomic group were compared with the entire 5,414 genomes, and Mann–Whitney P-values and fold-change were calculated.

## Data availability

Raw metagenomic sequencing data are available from the Sequence Read Archive (accession number will be released upon publication). By accessing the web server, www.mbiomenet.org/HRGM/, users can browse and download all genomes for 5,414 representative species, their annotations, and metadata, including geographical origin, taxonomy, genomic content, and genome statistics. The five classes of protein catalogs, 16S rRNA sequences, and SNVs are also provided with their functional annotation and taxonomic origin.

## Competing interests

The authors declare no competing interests.

## Author contributions

CYK and IL conceived this study. CYK and ML constructed the catalog and performed bioinformatics analysis. SY constructed the web server. KK, DY, and HRK organized the study cohorts and provided the fecal samples. IL supervised the project. CYK, ML, and IL wrote the manuscript.

## Acknowledgments

This research was supported by the National Research Foundation funded by the Ministry of Science and ICT (2018R1A5A2025079, 2018M3C9A5064709, 2019M3A9B6065192) to IL. We appreciate the assistance from the KOBIC Research Support Program.

## Notes

https://www.mbiomenet.org/HRGM/

### Competing Interest Statement

The authors have declared no competing interest.

