## Supplementary Figures for "Human reference gut microbiome comprising 5,414 prokaryotic species, including newly assembled genomes from under-represented Asian metagenomes"

**Supplementary Fig. 1-11**

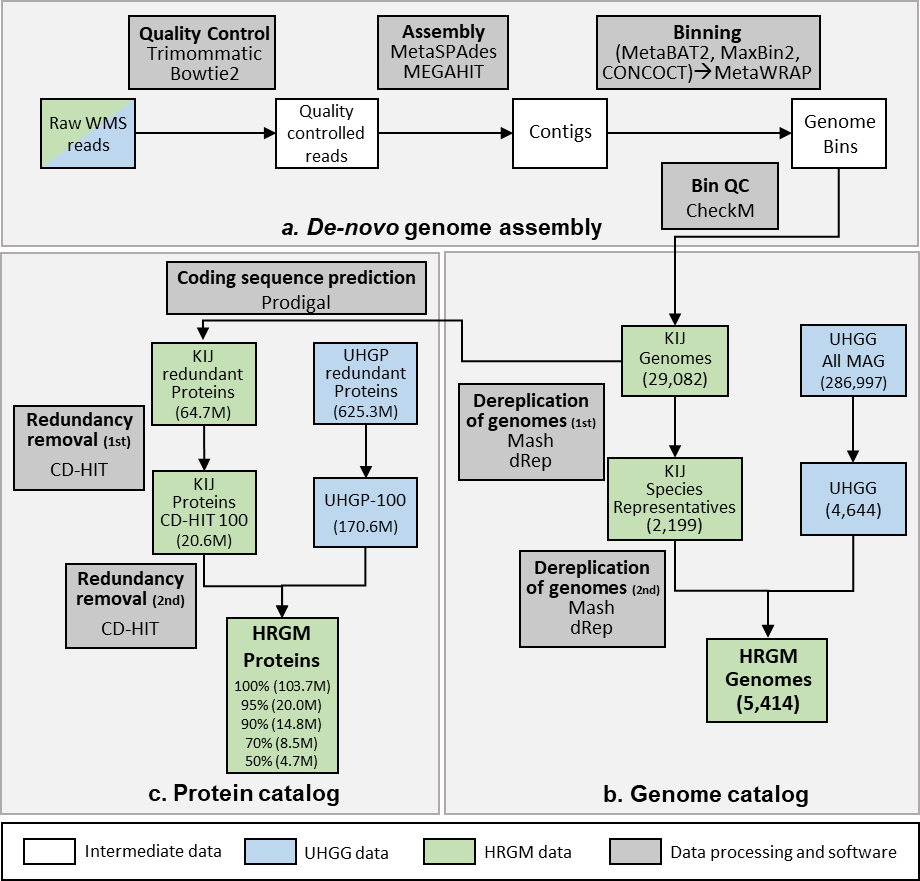

**Supplementary Fig. 1 | Overview of computational pipeline for cataloging genomes and proteins from whole metagenomic shotgun sequencing data.**

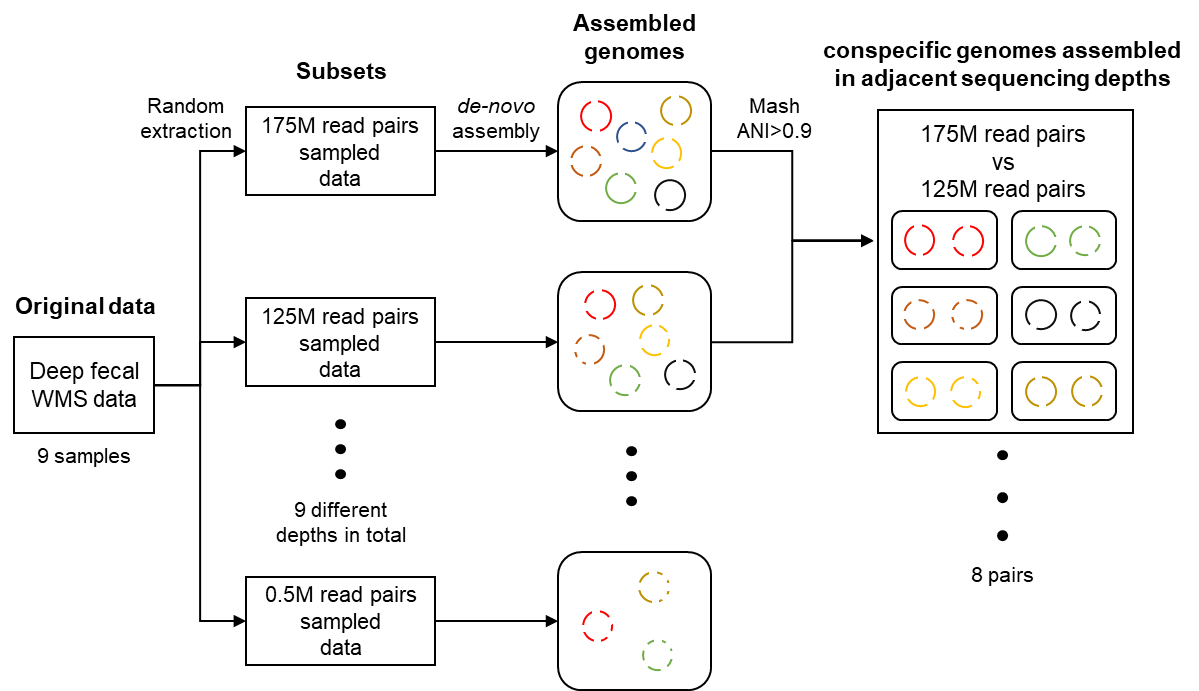

**Supplementary Fig. 2 | Overview of generating simulated data sets for assessing effect of sequencing depth on *de novo* genome assembly.** Flow diagram that describes methods of assessing the effect of sequencing depth on genome assembly with simulated data. We performed a *de novo* genome assembly on 81 simulated samples (9 depth-levels from 9 original samples). Genomes with the same color indicate that the genomes originated from the same species.

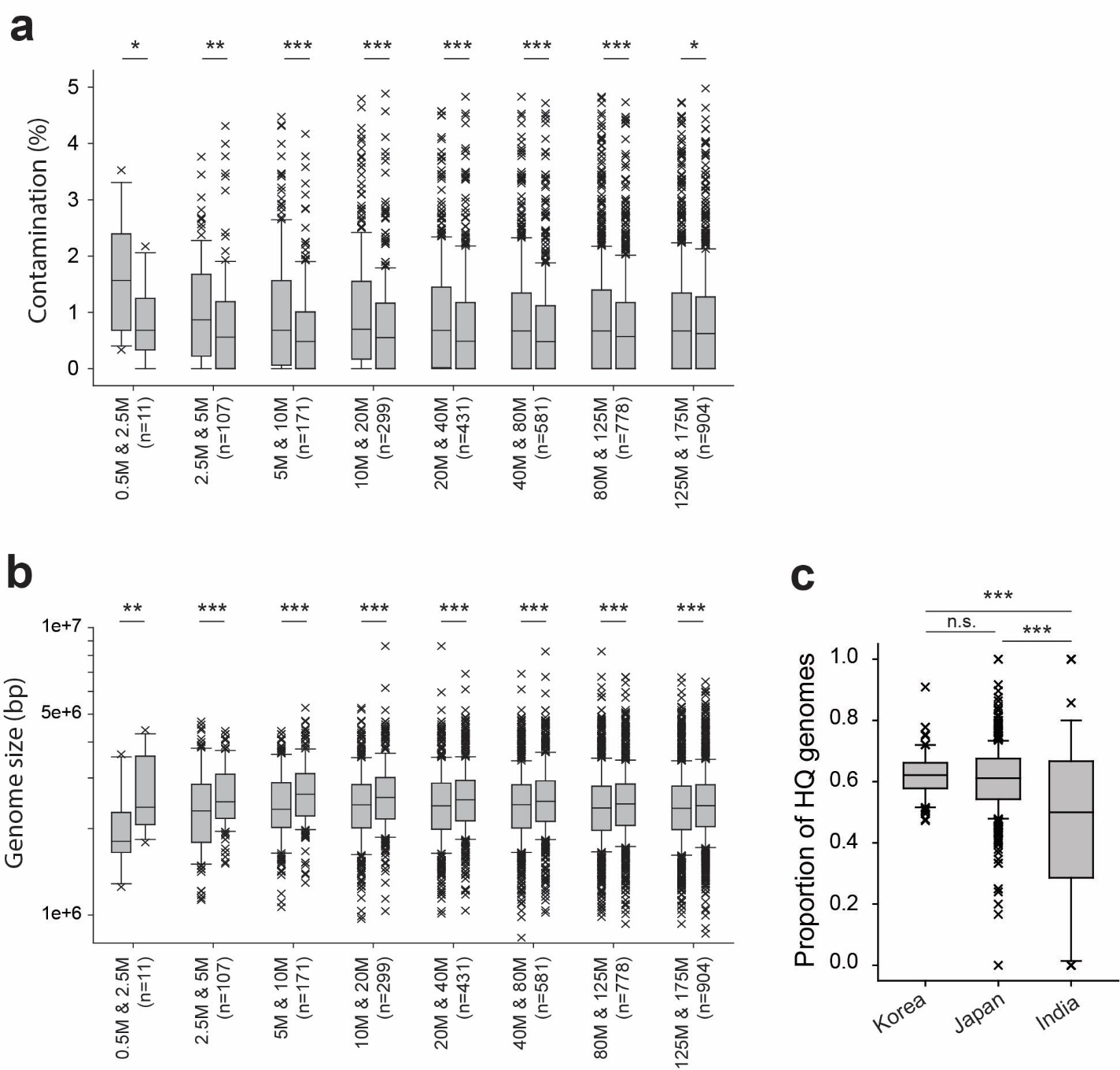

**Supplementary Fig. 3 | Quality of assembled genomes increases as sequencing depth increases.** The quality of the same genome assembled from different sequencing depth was evaluated by **a.** contamination, **b.** size of the genome. **c.** The proportion of high-quality genomes assembled from Korea, Japan, and India. *P*-values were evaluated by the two-sided Mann-Whitney U test. (n.s: *P* > 0.05, *: *P* < 0.05, **: *P* < 0.01, ***: *P* < 0.001)

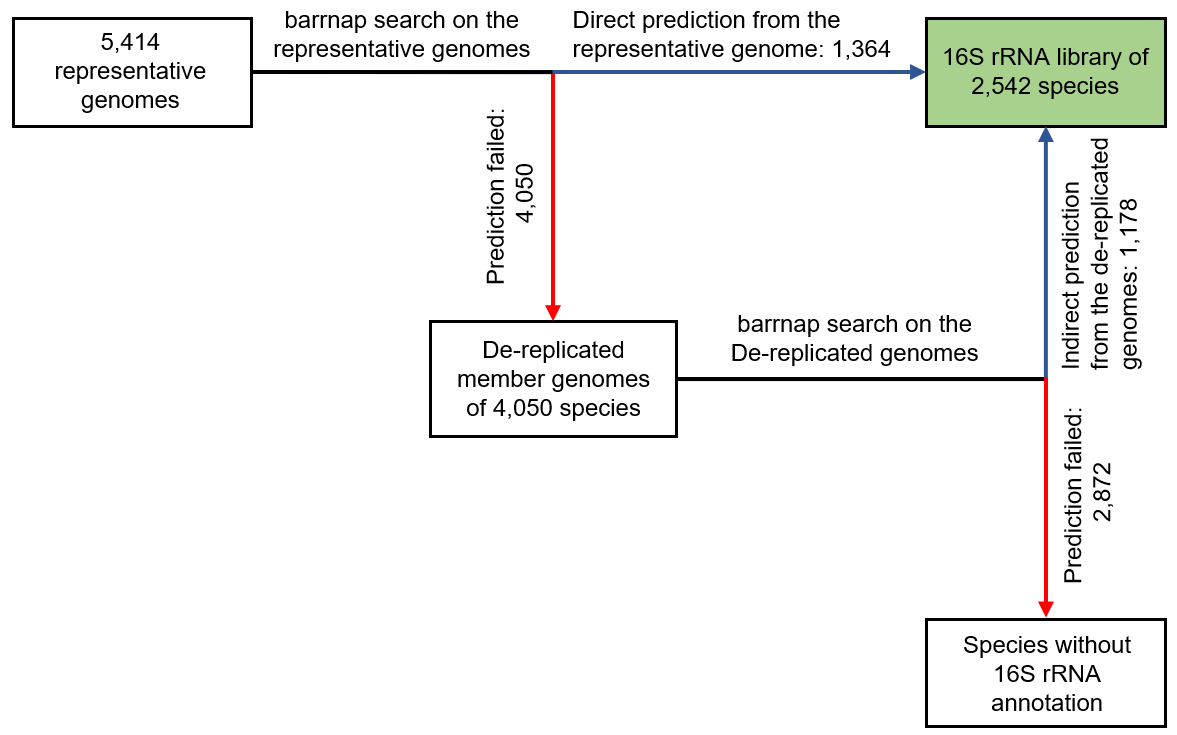

**Supplementary Fig. 4 | Computational pipeline for predicting 16S rRNA sequence region.** 16S rRNA regions were predicted with barrnap from representative genomes, and from de-replicated member genomes when failed to predict from representative genomes. Blue arrows represent the success of barrnap prediction, and red arrows represent the failed prediction.

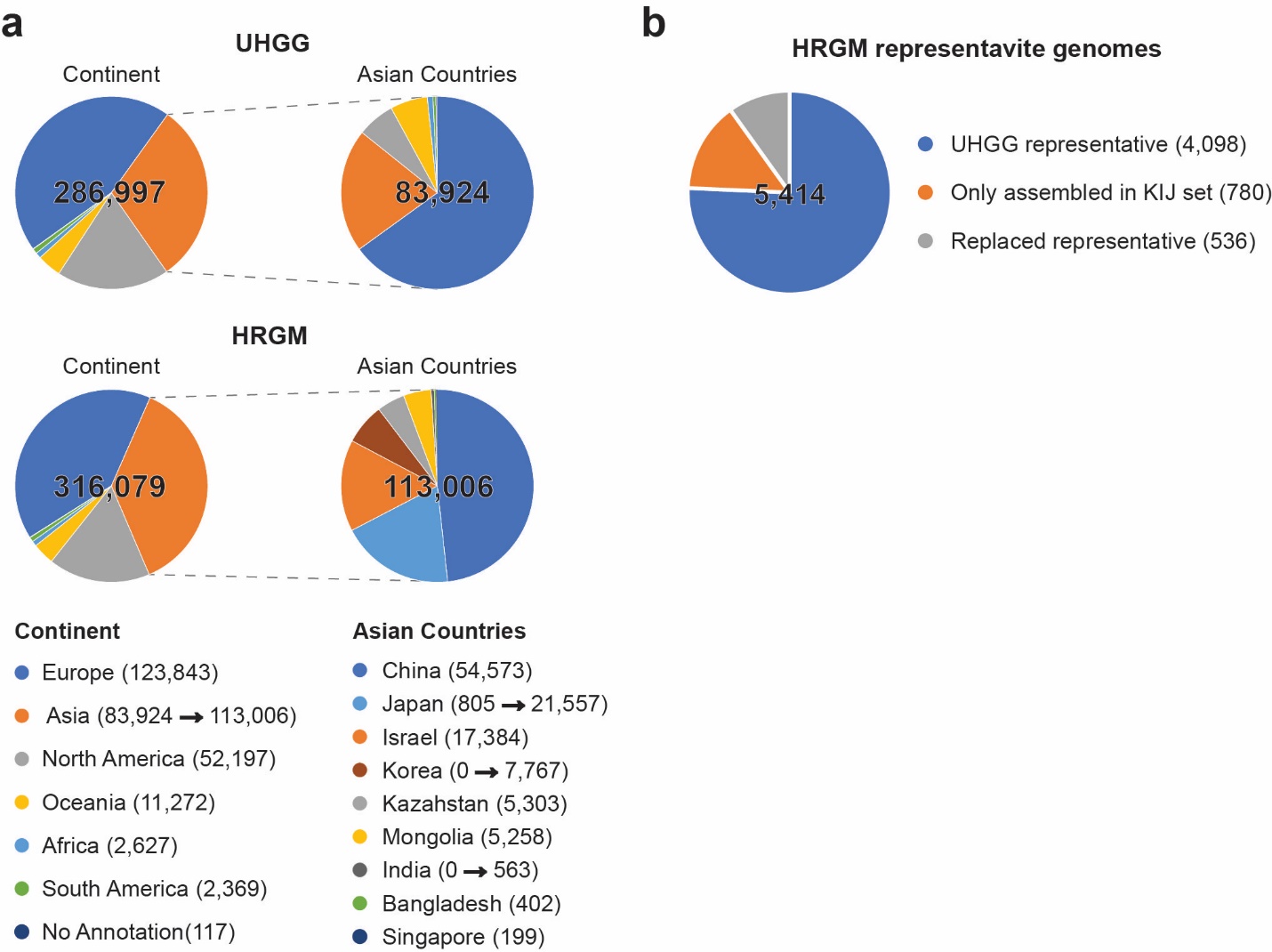

**Supplementary Fig. 5 | Comparison between UHGG and HRGM a,** Pie charts representing the number of MAGs by continents (left) and Asian countries (right) of UHGG (upper) and HRGM (lower) genome catalog. The exact number of MAGs for each continent and country is represented in the below legend. Arrows represent the change of MAGs from UHGG to HRGM. **b,** The number of representative genomes by their originated datasets.

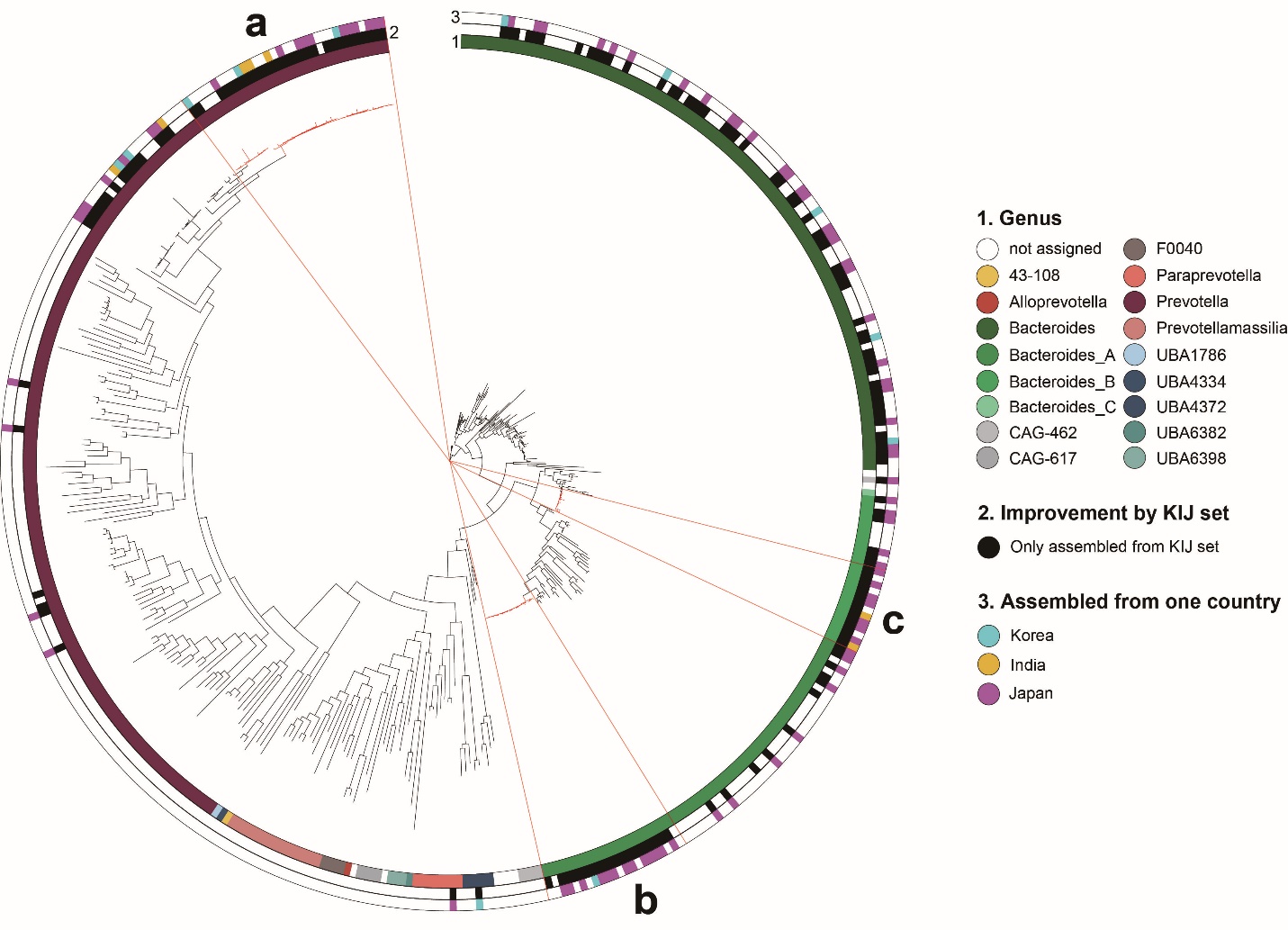

**Supplementary Fig. 6 | Phylogenetic tree of Bacteroidaceae family**. 410 branches in this family include many new MAGs from KIJ set. Three color strips indicate (1) genus annotated by GTDB-Tk (2) Improvement by KIJ samples (3) Assembly status for each country. There are three marked regions where phylogenetic tree branches are overdispersed. **a,** The region belongs to Prevotella genus and includes 30 genomes annotated as Prevotella copri. **b,** The region belongs to Bacteroides genus and includes 22 genomes annotated as Bacteroides plebeius. **c,** The region belongs to Bacteroides genus and includes 12 genomes annotated as Bacteroides vulgatus.

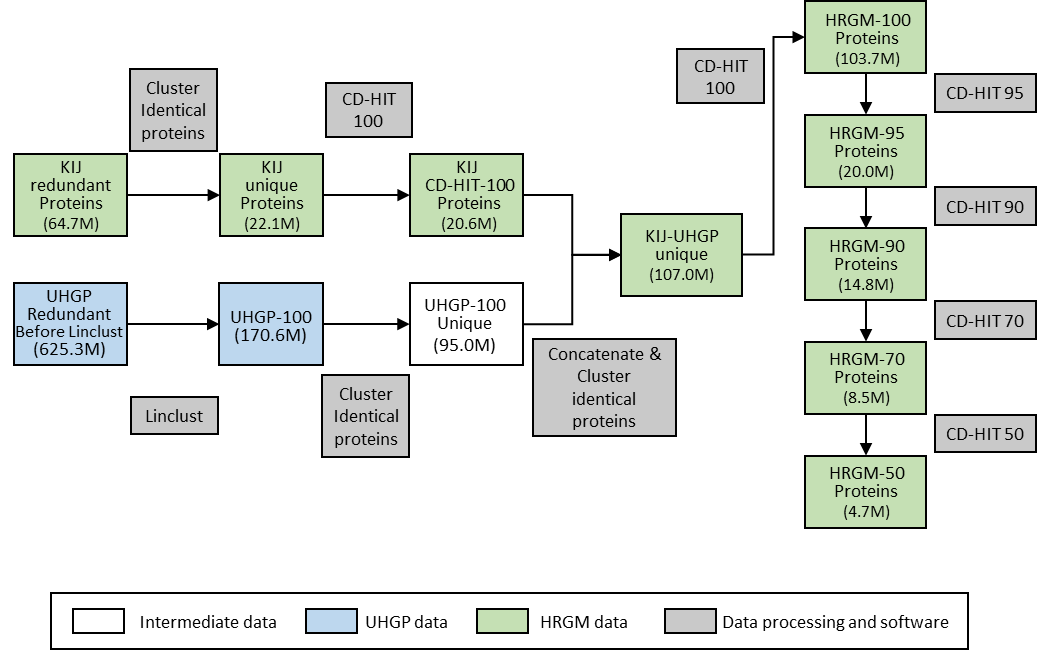

**Supplementary Fig. 7 | Overview of computational pipeline for cataloging nonredundant proteins.** The number in the parentheses indicates the number of proteins after each step. Information of proteins for the green boxes are freely available.

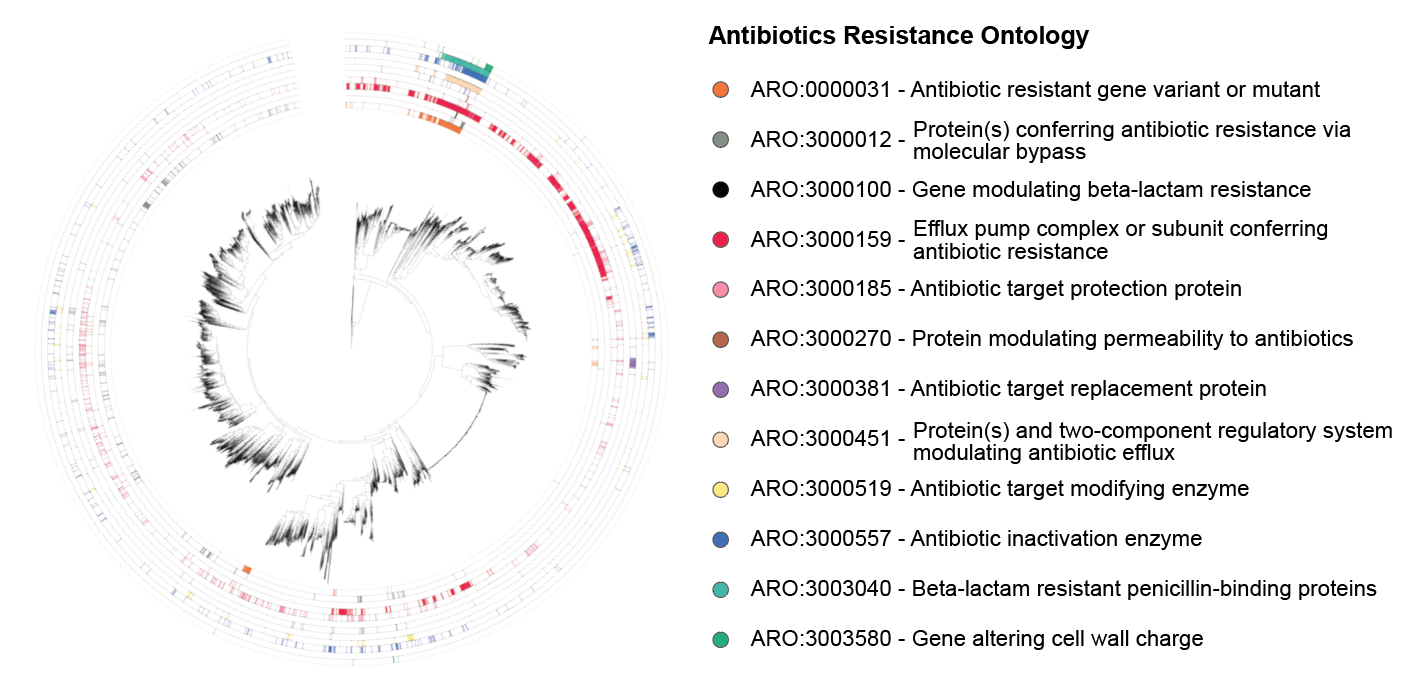

**Supplementary Fig. 8 | The landscape of antibiotics resistance ontology of human gut prokaryotic species.** Genomes having antibiotic resistance gene(s) are annotated according to ARO (antibiotic Resistance Ontology) terms. 12 labels are third level ARO terms which are annotated by at least one genome for each of the 5,414 HRGM representatives.

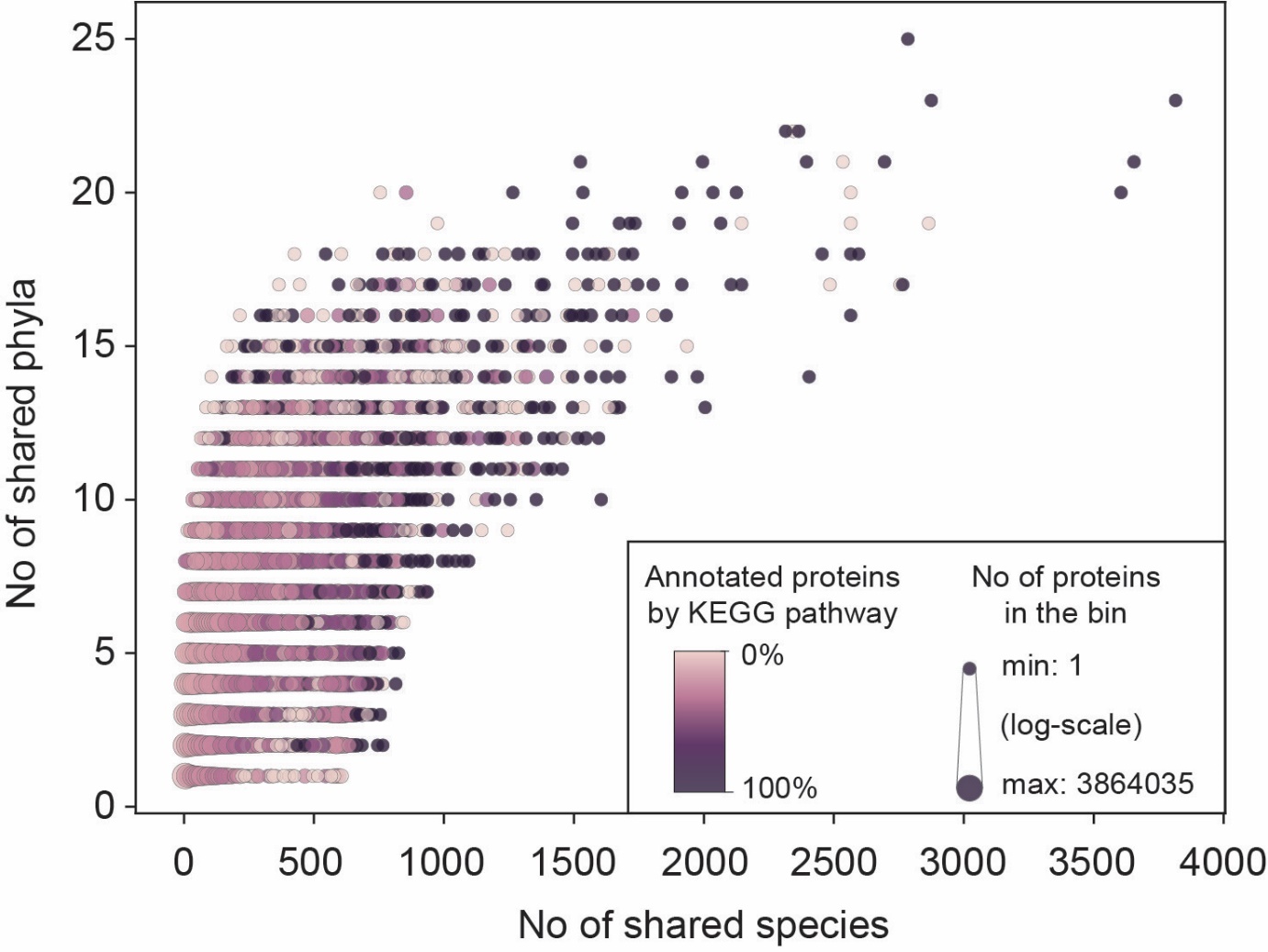

**Supplementary Fig. 9 | Gut microbial proteins that are shared by many species tend to be functionally annotated.** Binned scatter plot that represents the annotation rate of each protein bin. Proteins of the HRGM Protein-50 catalog were sorted by the number of their encoding species (X-axis) and annotation rates were measured for every bin of 10 proteins. Bright color indicates the proteins poorly annotated by the KEGG pathway. The size of data point represents the number of proteins in each bin.

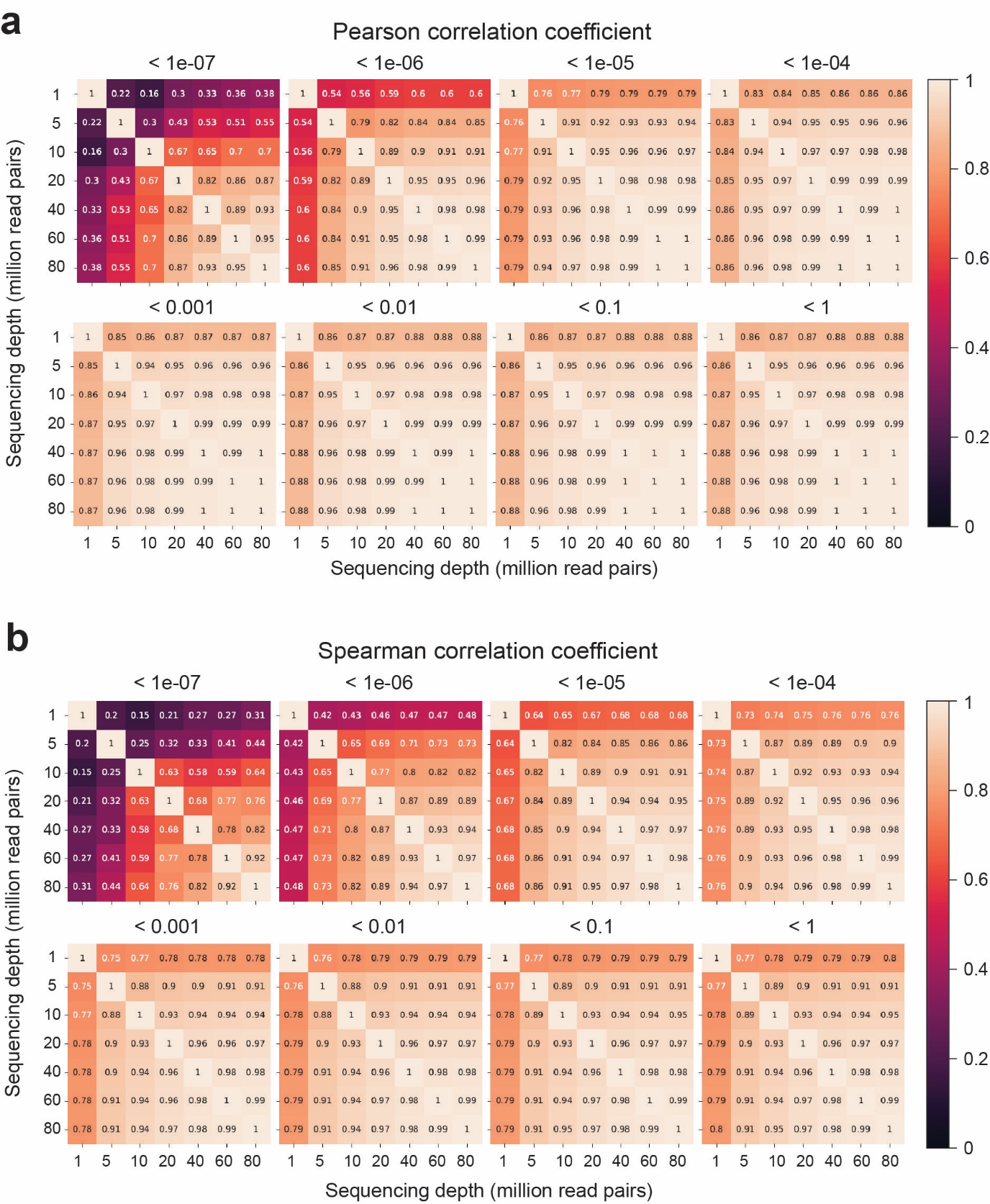

**Supplemental Fig. 10 | The effect of sequencing depth on taxonomic profiles based on WMS data**. **a**, Pearson correlation coefficient and **b**, Spearman correlation coefficient of the taxonomic profiles between the various sequencing depths (x- and y- axis) for the given mean relative abundance thresholds (title of each heatmap).

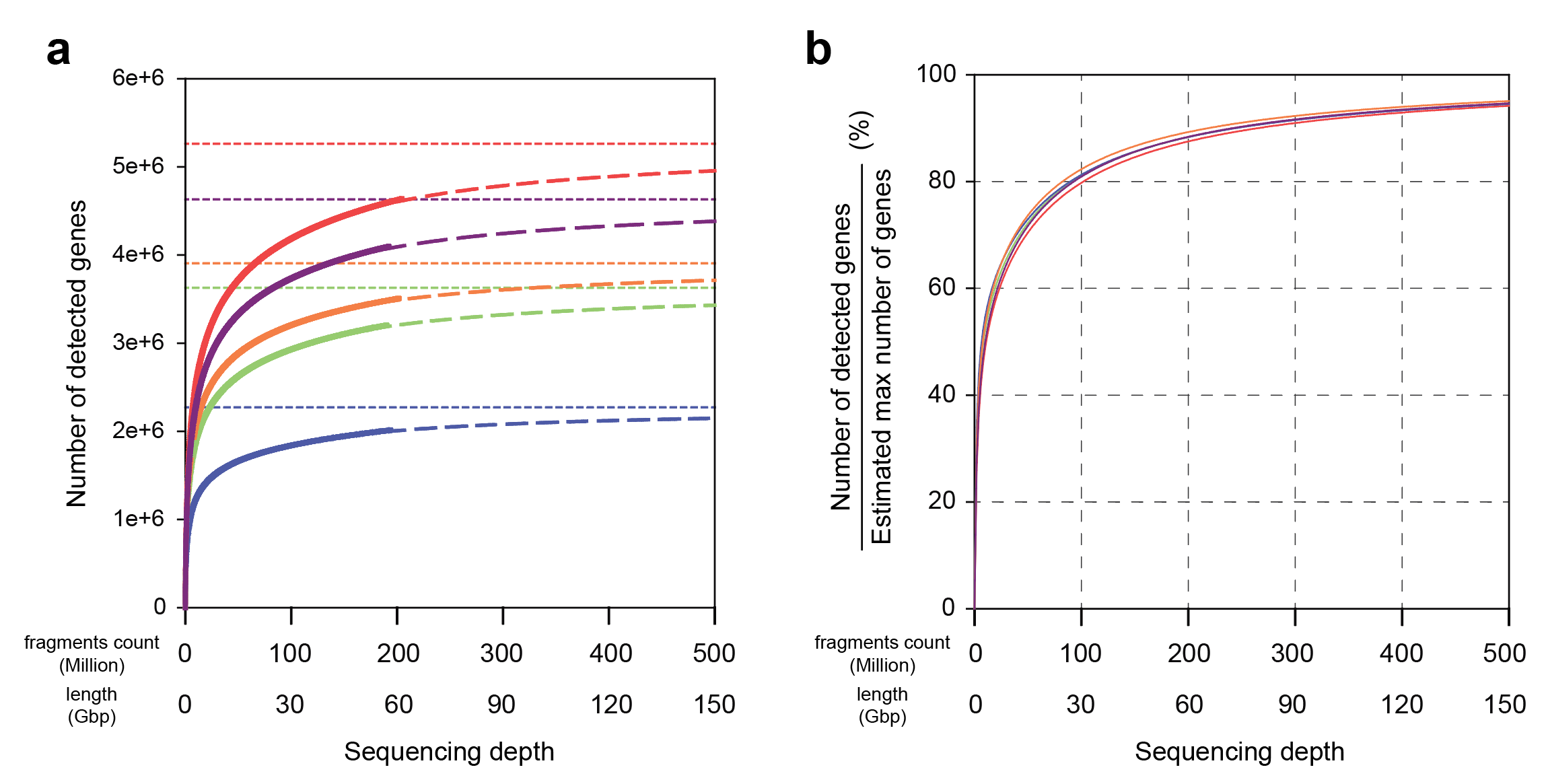

**Supplementary Fig 11 |** **Relationship between sequencing depth and the number of detected coding genes.** **a,** The number of detected gut microbial coding genes grows as fecal WMS sequencing depth increase. Different colors represent different samples. Solid curves denote the actual detected number of genes, and dashed curves show the regression curves by two-site saturation model. Horizontal dashed lines indicate the estimated max number of coding genes for each sample. **b,** The number of detected coding genes was normalized by the estimated maximum number of genes for given sequencing depth.
